# Two subsets of human marginal zone B cells resolved by global analysis of lymphoid tissues and blood

**DOI:** 10.1101/2021.03.19.436131

**Authors:** Jacqueline H.Y. Siu, Michael J. Pitcher, Thomas J. Tull, William Guesdon, Lucia Montorsi, Charles W. Armitage, Krishnaa T. Mahbubani, Richard Ellis, Pawan Dhami, Katrina Todd, Ulrich D. Kadolsky, Michelle Kleeman, David P. D’Cruz, Kourosh Saeb-Parsy, Mats Bemark, Gavin J. Pettigrew, Jo Spencer

**Affiliations:** Department of Surgery, University of Cambridge and NIHR Cambridge Biomedical Research Centre, Cambridge, CB2 0QQ, UK; School of Immunology and Microbial Sciences, King’s College London, Guy’s Campus, London, SE1 9RT, UK; School of Cancer Sciences, King’s College London, Guy’s Campus, London, UK; Cancer Systems Biology Laboratory, Francis Crick Institute, London, UK; Biomedical Research Centre, Guy’s and St. Thomas’ NHS Trust, London, SE1 9RT, UK; Department of Microbiology and Immunology, Institute of Biomedicine, Sahlgrenska Academy, University of Gothenburg, SE 405 30 Gothenburg, Sweden; Department of Clinical Immunology and Transfusion Medicine, Region Västra Götaland, Sahlgrenska University Hospital, Gothenburg, Sweden

## Abstract

B cells generate antibodies that are essential for immune protection. Major events driving B cell responses occur in lymphoid tissues, which guide antigen acquisition and support cellular interactions, yet complexities of B cell subsets in human lymphoid tissues are poorly understood. Here we perform undirected, global profiling of B cells in matched human lymphoid tissues from deceased transplant organ donors and tracked dissemination of B cell clones. In addition to identifying unanticipated features of tissue-based B cell differentiation, we resolve two clonally independent subsets of marginal zone B cells that differ in cell surface and transcriptomic profiles, tendency to disseminate, distribution bias within splenic marginal zone microenvironment and immunoglobulin repertoire diversity and hypermutation frequency. Each subset is represented in spleen, gut-associated lymphoid tissue, mesenteric lymph node, and also blood. Thus, we provide clarity and diffuse controversy surrounding human MZB - the ‘elephant in the room’ of human B cell biology.

## Introduction

B cells maintain health by generating affinity-matured and innate-like antibody responses, and as immune regulators that present antigen and secrete cytokines. B cell responses occur in lymphoid tissues that differ in fundamental microarchitecture and mechanisms of antigen acquisition (Junt et al., 2008). For example, gut-associated lymphoid tissue (GALT) in the Peyer’s patches and appendix chronically sample predominantly particulate antigens from the gut lumen via a specialised follicle associated epithelium (FAE). This results in sustained germinal centre (GC) responses (Yamanaka et al., 2003). In contrast, lymph nodes (LN) receive soluble or complexed antigens via afferent lymphatics (Drayson and Ford, 1984). Apart from mesenteric lymph nodes (mLN) that are also associated with modulating immunity on mucosal surfaces and that are chronically stimulated by gut-derived antigen, lymph nodes in healthy humans tend to be quiescent and lack GC (Houston et al., 2016; Pabst and Mowat, 2012). Similarly, the spleen, which receives antigens delivered from the blood, is also generally immunologically quiescent in a healthy human (Wilkins and Wright, 2000).

Despite the intrinsic importance of tissue microanatomy and antigen acquisition for B cell responses, there has been a tendency to depend on studies of blood by flow cytometry, which is highly directed, to establish definitions of human B cell subsets and changes within them in disease (Sanz et al., 2019). Blood, which only accommodates a small fraction of all lymphocytes, contains a conglomerate population of cells with different migratory biases as demonstrated by differences in expression of receptors for tissue site-associated endothelial ligands (Vossenkämper et al., 2013). Some subsets in blood, such as the circulating marginal zone B (MZB) cells, are named according to splenic microanatomy by shared phenotype, but the extent to which they are truly analogous is unclear (Kibler et al., 2021; Mebius et al., 2004; Weill et al., 2009; Weller et al., 2004). In addition, whilst the connectivity between tissues that are bridged by migrating cells has been identified by repertoire analysis and by some deep phenotypic comparisons, the extent to which different B cell subsets share clonal relatives and whether they remain tissue resident or migrate between tissue sites is unknown (Magri et al., 2017; Nair et al., 2016; Weisel et al., 2020; Zhao et al., 2018).

Here we determine the characteristics and relative distribution of human B cell subsets in matched samples of spleen, mLN and appendiceal GALT using deep phenotyping, single cell transcriptomics and clonotype analysis. The strategy was to use undirected methods to group cells and then to infer identities of the populations thus identified from published work. We observe several subsets of activated B cells in tissues that are absent in blood and uncover evidence for tissue-based maturation from transitional (TS) to naive B cells. We then focus our attention on two subsets of MZB cells in tissues and blood and that we designate MZB-1 and MZB-2, since the nature of such cells is a controversial area needing urgent clarification (Nemazee, 2021). MZB-1 and -2 cells differ in transcriptomic profiles, levels of repertoire diversity, hypermutation frequency, their tendency to disseminate, and in their distribution bias within the MZ. Our observations bring features of MZB cells into sharp focus within undirected multidimensional representations of human B cells in tissues and blood. The identification of distinct MZB-1 and MZB-2 subsets reconciles previous key, yet apparently conflicting observations relating to human MZB cells (Descatoire et al., 2014; Jenks et al., 2018; Johnson et al., 2020; Kibler et al., 2021; Nemazee, 2021; Tipton et al., 2015; Tull et al., 2021; Vossenkämper et al., 2013; Zhao et al., 2018), and provides needed clarity in an area that has been in turmoil for many years.

## Results

### Regional variation in abundance of B cell phenotypic variants in tissues

Mass cytometry using a panel of 28 antibodies was used to investigate regional variation in B cells in human lymphoid tissues (Supplementary Table 1). We analysed 8 matched samples of appendix, mLN, and spleen from deceased transplant organ donors with no known disease (Fig. S1 A). Following quality control (Fig. S1, B and C), data were batch normalised (Fig. S1, D and E) (Finck et al., 2013; Lun et al., 2017). We then made a 3-way undirected comparison of the differential abundance of B cells in the tissues according to both position on the UMAP and intensity of marker expression, using the Cydar package (Lun et al., 2017). Rather than clustering, ‘hyperspheres’ generated by Cydar that represent the cells with both similar position on the UMAP and similar median marker intensity were analysed (McInnes et al., 2018). Only hyperspheres containing 100 cells or more were used for downstream analysis (Fig. S1 F).

The expression of key markers including CD27, IgD, IgM, IgA and CCR7 by hyperspheres was visualised on the UMAP (Fig. 1 A). The cell count for individual hyperspheres on the UMAP demonstrated similar cell densities, with slightly higher number of cells within CD27^-^ and CD27^+^IgM^high^ hyperspheres than those representing class switched-cells (Fig. S1 G). Cell counts of hyperspheres identified similar distribution between donors within spleen and appendix samples, whilst mLN samples were more variable falling between the other organs or overlapping with GALT (Fig. S1 H). A three-way comparison of differential abundance of hyperspheres between tissues was performed. Hyperspheres with significantly different abundances between tissues were predominantly CD27^+^IgM^+^, suggesting that the predominant inter-tissue variability is among unswitched, antigen-experienced cells (Fig. 1, A and B).

**Fig. 1:**
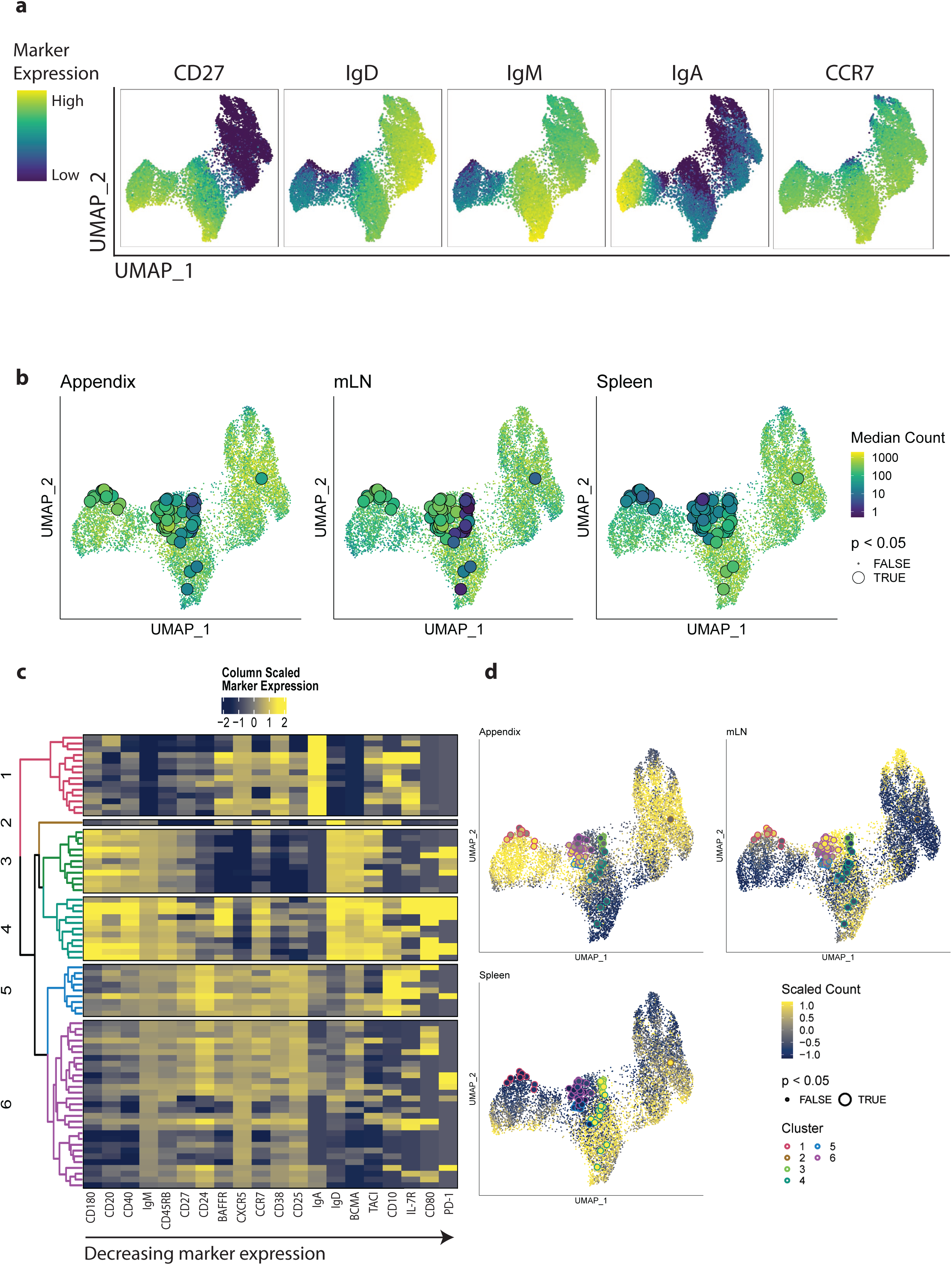
Differentially abundant B cell subpopulations between human lymphoid tissues. **(A).** UMAP plot of the median positions of hyperspheres for CD19^+^ B cells in concatenated human appendix, mLN, and spleen. Each point represents a hypersphere coloured by the median intensity of selected markers (bounded by the 5th and 95th percentiles of the intensities across all cells) for that hypersphere. **(B).** UMAP coloured by cell counts within each hypersphere for each tissue. The larger points represent the significantly differentially abundant hyperspheres detected at a spatial FDR of 5%. **(C).** Heatmap of markers expressed in hyperspheres with significantly difference abundances, scaled by column to highlight differences between clusters. Hyperspheres were clustered by hierarchical clustering and k-means with coloured dendrogram to identify clusters. **(D).** UMAP coloured by scaled hypersphere count across the three tissues and outline coloured by cluster as in **(C).**

We next compared the phenotypic features of cells in the hyperspheres that differed significantly in abundance between tissues, using hierarchical clustering and k-means. This identified 5 major CD27^+^ clusters of hyperspheres and one CD27^-^ cluster (Fig. 1 C). The hyperspheres in each cluster were colour coded according to the dendrogram and located on the UMAP for each tissue (Fig. 1 D).

The single hypersphere in cluster 2, that had significantly higher abundance in spleen, contained cells expressing IgM, IgD, CCR7 and CD10, although it did not express high levels of characteristic TS B cell markers CD38 or CD24. This small cluster was most closely linked in the dendrogram to the two further clusters of hyperspheres (clusters 3 and 4) that were also both relatively more abundant in spleen, and that had the CD27^+^IgM^+^IgD^+^ phenotype of MZB cells. While both phenotypically resembling MZB, one subset had significantly higher expression of CCR7, CXCR5, BAFFR, CD24 and CD27 than the other. The remaining clusters of hyperspheres 1, 5 and 6, that were all most abundant in GALT and mLN, comprised IgA-expressing B cells including some CD10 expressing GC variants (cluster 1), IgM expressing GC cells (cluster 5) and IgM-only cells with the phenotype CD27^+^IgM^+^IgD^-^ (cluster 6) (Fig. 1, B-D).

### Regional B cell complexity dissected by single cell transcriptomics

To understand B cell variability in tissues in more depth, three representative sets of tissues from single individuals were selected for analysis by single cell RNA sequencing (scRNAseq) (Fig. S1 A). To facilitate identification of B cell subsets, sorted CD19^+^ cells were surface labelled with Total-Seq-C antibodies prior to capture on the 10x chromium controller (Fig. S2, A and B). Gene expression, antibody detection tag (ADT) and V(D)J libraries were then prepared according to the manufacturer’s instructions and sequenced. Following quality control and normalisation, data from all tissues and donors were integrated (Fig. S2, C-F). After checking that cells did not separate according to donor on a UMAP (Fig. S2 G), cell clusters were subsequently identified according to the expression of the 3000 variable genes. The expression of key genes and cell surface markers were visualised on the UMAP (Fig. 2 A).

**Fig. 2:**
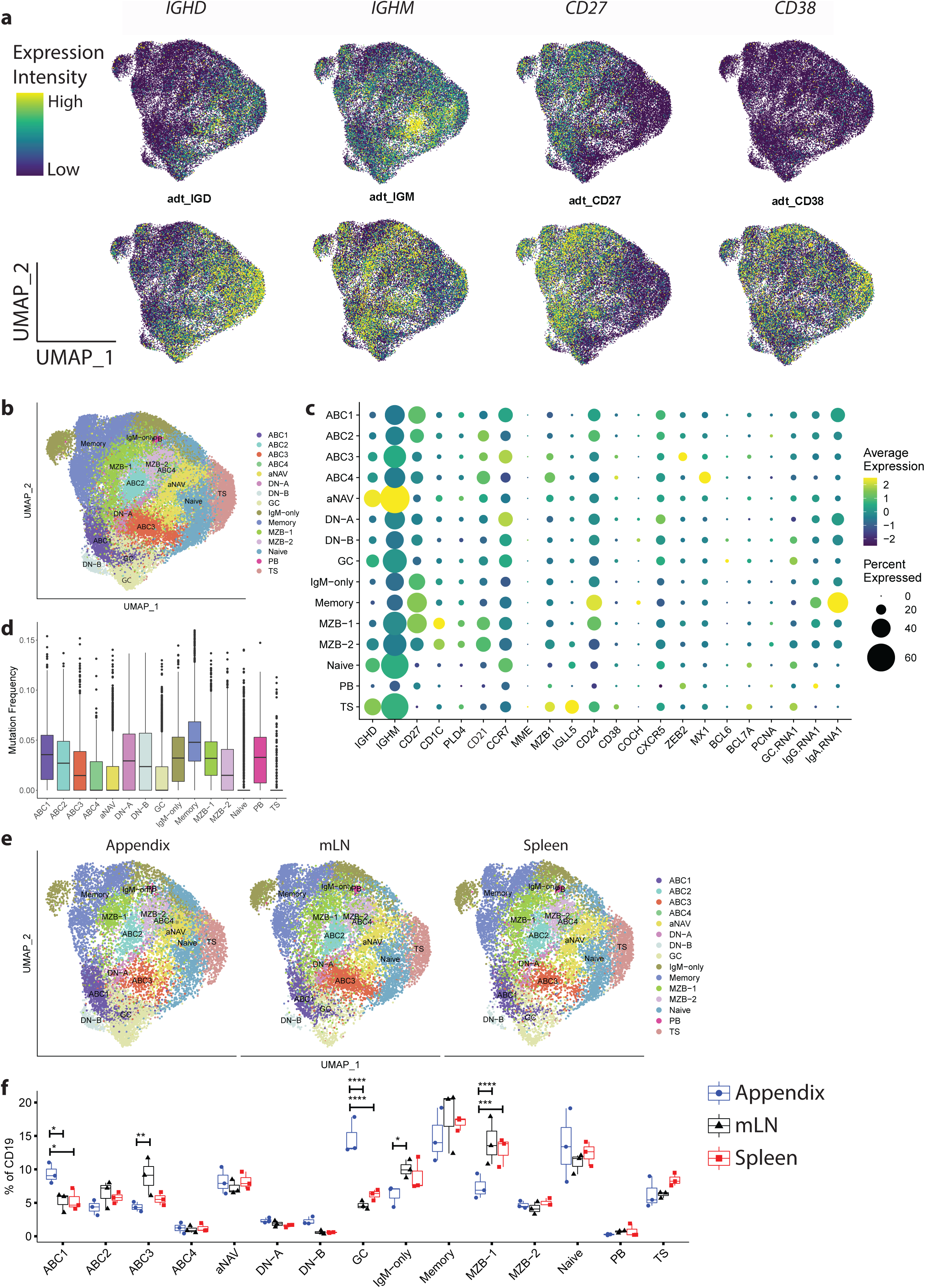
Overview of B cell composition of human lymphoid tissues from single cell transcriptome and BCR profiling. **(A).** UMAP visualisation of select mRNA transcripts (top) and ADT surface protein (bottom) of total CD19^+^ B cells. **(B).** UMAP visualisation of B cell composition in lymphoid tissues coloured by B cell subsets, annotated based on Seurat unsupervised clusters (ABC1-4, activated B cells 1-4; aNAV, activated naive; DN, double negative; GC, germinal centre B cells; MZB, marginal zone B cells; PB, plasmablast; TS, transitional B cells). **(C).** Dot plot illustrating marker gene expression for B cell subtypes. The following gene groups were used: GC.RNA.1 (*BCL6, BCL7A*); IgG.RNA.1 (*IGHG1, IGHG2, IGHG3, IGHG4*); IgA.RNA.1 (*IGHA1, IGHA2*). **(D).** Frequency of somatic mutations in *IGHV* genes used by B cells in each B cell subset. **(E).** UMAP visualisation of B cell subsets in each lymphoid tissue. **(F).** Relative proportion of B cell subsets in each lymphoid tissue. Statistics for differential abundance between matched tissues for each subset were assessed by estimated marginal means with Bonferroni correction for multiple comparisons. *p < 0.05, **p < 0.01, ***p < 0.001 and ****p < 0.0001.

Sixteen distinct clusters were identified and named by reference to transcript profile and cell surface phenotype (Fig. S3) and visualised on the UMAP (Fig. 2 B, and summarised in Fig. 2 C). Some clusters included cells that were easily recognisable according to known profiles of gene expression and cell surface phenotypes described before. These were TS, naive, activated naive (aNAV), GC, and class switched memory B cells. Markers that identified them are highlighted in Fig S3. The aNAV subset, that is a reported precursor to antibody secreting cells derived from the extrafollicular response, was differentiated from naive and TS B cell subsets by its high expression of *CD19* and *ITGAX (CD11c)*, and low expression of *CXCR5, CD24*, and *CD38* (Fig. S3 C) (Jenks et al., 2018; Tipton et al., 2015).

Four clusters of cells that had markers of activation but no classic markers of GC (*BCL6, BCL7a*), were identified and designated activated B cell (ABC) 1 to 4. Cells classified as ABC1 resembled GC B cells most closely, expressing *JUN, JUNB* and *FOS*. Cells classified as ABC2 had strong expression of the proliferation antigen *PCNA* but lacked other markers of activation. ABC3 was designated ‘activated’ because it expressed the long non-coding RNA *MALAT1* (Fig. S3). *MALAT1* is linked to class switching through its role in the alternative-non-homozygous end joining pathway (Kato et al., 2012). This cluster also expressed the *ZEB2* transcription factor associated with aNAV and the extrafollicular response(Sanz et al., 2019). ABC4 had a strong signature of interferon regulated genes suggesting prior activation via this route.

Two non-adjacent clusters containing relatively few cells were termed double negative (DN) because they lacked both CD27 and IgD (Fig. 2 A). However, they did not separate according to the classical features that discriminate between subsets of DN cells in blood (Fig. S3 D).

Two clusters of MZB defined by cell surface CD27^+^IgM^+^IgD^+^ phenotype and presence of *CD27, IGHM, IGHD, CD1C* and *PLD4* transcripts were identified (Fig. 2 A-C).

Frequencies of somatic hypermutation of *IGHV* genes (SHM) of cells in clusters were generally consistent with expected properties according to cluster definitions (Fig. 2 D). The extent of SHM was highest in memory B cells and lowest in naive and TS B cells. GC B cells had undergone relatively few rounds of SHM. Clusters ABC1-3 each had moderate SHM as had the MZB-1 subset, similar to IgM-only memory cells. Cells in MZB-2 had lower levels of SHM than MZB-1.

The relative abundance of the B cell subsets between tissues was compared (Fig. 2, E and F). GC cells and ABC1 cells were both more abundant in the appendix than mLN or spleen. IgM-only B cells were more abundant in mLN than appendix as were ABC3 cells. The MZB-1 subset was proportionally underrepresented in appendix compared to other tissues. MZB-2 cells were lower in frequency than MZB-1 cells, but not significantly more or less proportionately abundant at any tissue site.

### Developmental relationships between B cell subsets in tissues

We investigated the relationships between cells in subsets using BCR and RNA velocity analysis (Fig. S4 A). Using the standard Immcantation pipeline for analysis of adaptive immune receptor repertoire in heavy chain genes, clones were identified from the hamming distance between each sequence and its nearest neighbour (Fig. S4 B) (Gupta et al., 2015). Clonal abundance (number of cells in clones as a proportion of the entire repertoire) was greater in the spleen than other tissues (Fig. S5 A). Of the antibody isotypes IgG tended to have greater clonal abundance (Fig. S5 A). Clonal diversity between isotypes, tissues, and subsets were visualised using a diversity profile curve; spleen and IgG had lower biodiversity compared to other tissues and isotypes (Fig. S5 B). There was no significant CDR3 length difference across isotypes, tissues, or subsets (Fig. S5 C).

We compared the tendency for subsets to contain migratory clones by plotting the ratio of clones with members observed in only one tissue to those observed in multiple tissue sites for each subset (Fig. 3 A). The subsets that contained more clones found in multiple tissues than in a single tissue site in all donors were IgM-only cells, MZB-1, memory cells and the ABC3 subset. Notably, clones with members designated as GC tended to remain local.

**Fig. 3:**
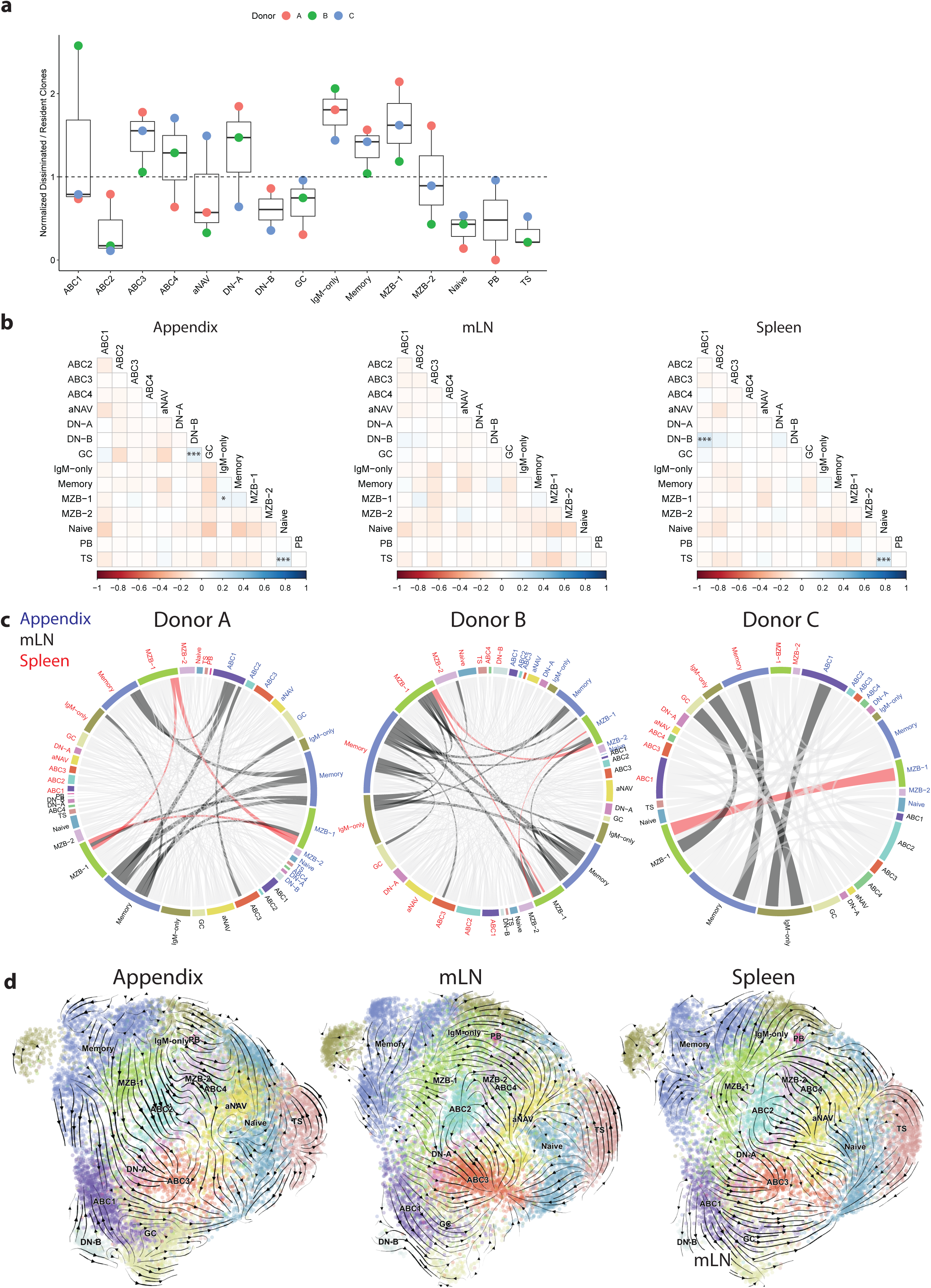
Clonal relationship and dissemination of B cell subsets within and between tissues. **(A).** Proportion of clones observed in 2 or more tissues (disseminated) compared to clones in one tissue only (resident) associated with each B cell subset, normalised by donor. Values above the dotted line indicate more dissemination and values below the dotted line indicate more resident clones for the subset. **(B)**. Clonal relatedness between B cell subsets within each tissue illustrated in a correlation plot. Statistically significant tendencies for clonal relatedness were identified using Spearman correlation along with pairwise p-values that were then corrected using Holm’s method. **(C).** Circos plots showing the clonal relationships between B cell subsets in the appendix (blue text), mLN (black text), and spleen (red text) for Donor A (left), B (middle), C (right). Clonally related sequences across tissue-subsets are connected by lines, with the top 5% most frequent connections coloured in dark grey and with sequences spanning MZB-1 across tissues coloured in red. All other connections are coloured light grey. **(D)**. RNA velocities of B cells for each tissue shown on the UMAP plot in Fig. 2 B. *p < 0.05, **p < 0.01, ***p < 0.001.

We used a correlation matrix to evaluate the frequency of clone sharing between different B cell subsets that indicates a developmental relationship between subsets within tissue sites (Fig. 3 B). Significant clone sharing was observed between TS and naive B cells in both appendix and spleen. Importantly, clone members were only observed locally within the same tissue and not shared between different tissues. When both heavy and light chain rearrangements were captured, these were shared by clone members. Thus, this indicates proliferation once heavy and light chain genes have rearranged, rather than proliferation at the pre-B stage, and suggests that TS mature to naive B cells in GALT and spleen. The lack of shared clones between tissues indicates that clones become diluted once they leave the tissue site. The existence of such local clones involving immature cells in tissues is consistent with a low level of proliferation at this stage (Vossenkämper et al., 2013).

In the appendix, MZB-1 and IgM-only B cell subsets were significantly clonally related, as were GC cells and DN-B cells, that locate adjacent to each other on the UMAP plot. In spleen, significant clonal relationships were observed between the DN-B and ABC1 subsets, also proximal in the UMAP. No significant clonal relationships between subsets were observed in mLN (Fig. 3 B).

Circos plots were used to visualise clone sharing between subsets across tissues. The clones most likely spanning tissues in all three donors were those containing MZB-1 and also IgM-only B cells, though to a lesser degree (Fig. 3 C).

The analysis of developmental trajectories by RNA velocity largely supported the findings above (Fig. 3 D) (Bergen et al., 2020). Transcriptomic progression from TS to naive B cells was observed in GALT and spleen. Developmental connections between GC, ABC1 and DN-B were apparent. There was evidence of memory B cell movement towards GC in GALT (Bunker et al., 2017). Consistent with Fig. 3 B, MZB-1 and MZB-2 showed no apparent connection (Fig. 3 D).

The cluster ABC3 characterised by the expression of *MALAT1* and *ZEB2* appeared to be a common ‘destination’ for developmental trajectories, despite no evidence on the repertoire analysis of developmental associations with other subsets. This cluster possibly represents a transcriptomic state, rather than a stage in progression of specific subsets.

### Human MZB subsets

The above analyses indicate the presence in humans of two subsets of MZB that differ in their: phenotype (Fig. 1 C); SHM (Fig. 1 D); transcriptome (Fig. 2, B and C); their development relationship to other clusters (Fig. 3 B); and their tendency for clonal dissemination (Fig. 3, A and C). No evidence of clonal links between them was observed (Fig. 3 B).

Differentially expressed genes between MZB-1 and MZB-2 subsets are illustrated in a volcano plot (Fig. 4 A). Genes selectively expressed by MZB-2 cells tended to be associated with cellular activation such as the HLA alleles, *CD83* and *MIF (Krzyzak et al., 2016; Starlets et al., 2006)*. In addition, MZB-2 cells selectively expressed the RNA helicase *DDX21*, which has been implicated in recognition of viral RNA (Chen et al., 2014). Interestingly, *CD83* is consistently more abundantly expressed in the appendix in both MZB-1 and MZB-2 subsets. The preferential expression of *CD37* by MZB-1 cells is consistent with its observed higher motility (Fig. 4, A and B) (Yeung et al., 2018).

**Fig. 4:**
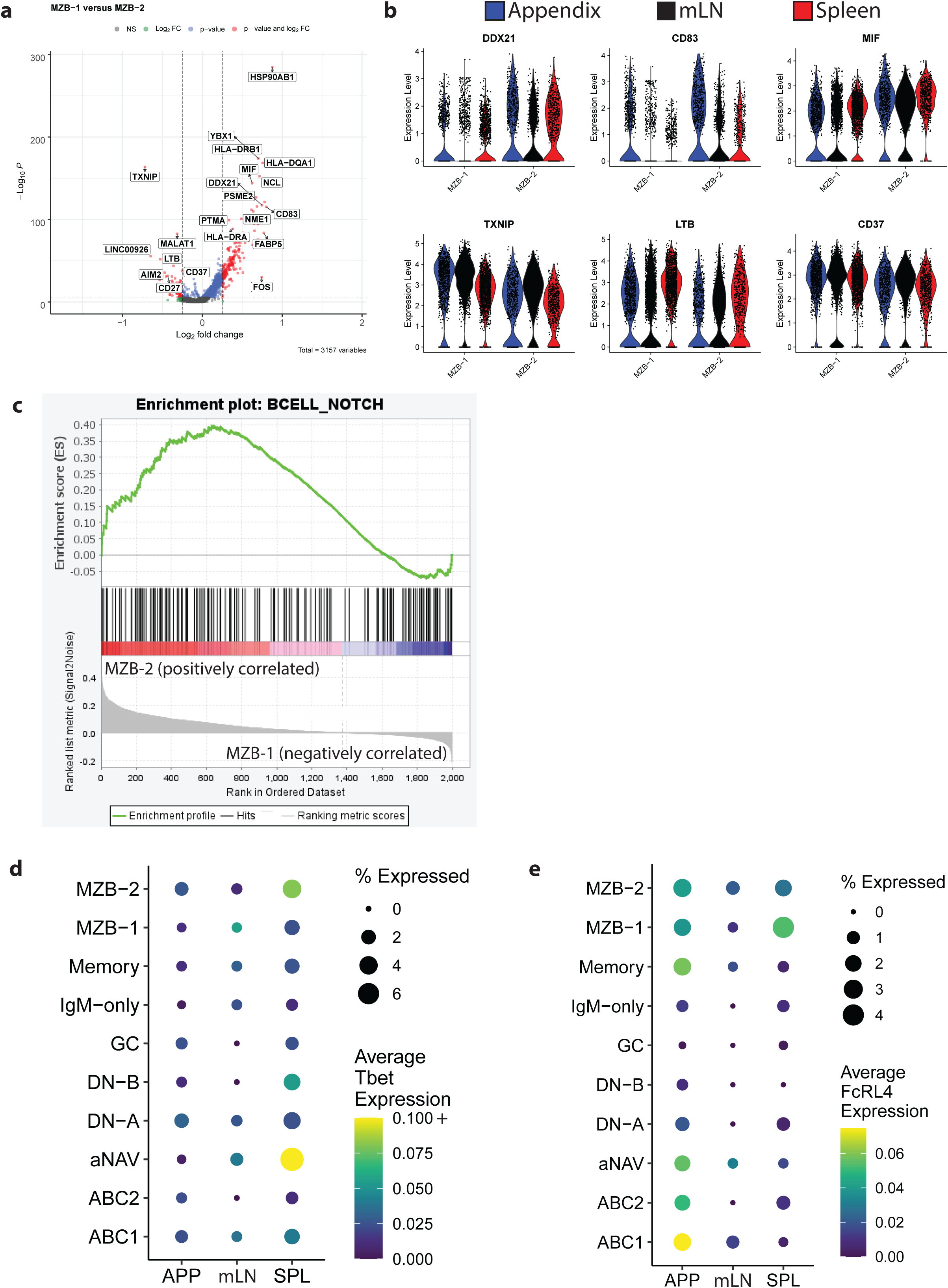
Transcriptomic differences between MZB-1 and MZB-2 B cell subsets. **(A).** Volcano plot comparing differentially expressed genes between MZB-1 and MZB-2. Cut offs for significant p value is 10^-6^, and log_2_FC is 0.25. Red dots highlight the significantly differentially expressed genes. P values were calculated using Wilcoxon test with Bonferroni correction for multiple comparisons. Due to space constraints, only select markers are labelled. **(B).** Violin plots demonstrating gene expression of *DDX21, CD83, MIF, TXNIP, LTB,* and *CD37* in MZB-1 and MZB-2 B cells across the three tissues. **(C).** Enrichment plot of the GSEA analysis using NOTCH gene sets from ImmuneSigDB between MZB-1 and MZB-2 cells, corrected for dataset background. Gene set is significantly enriched in the MZB-2 subset at nominal p value < 1%. **(D).** Dot plot illustrating the expression of the gene that encodes for Tbet transcription factor, *TBX21* for select B cell subtypes. **(E).** Dot plot illustrating *FcRL4* gene expression for select B cell subtypes.

MZB development in mice and humans involves ligation of NOTCH2 on B cells by delta like-1 (Descatoire et al., 2014; Saito et al., 2003). We therefore sought evidence of enrichment of genes in the NOTCH pathway in the MZB subsets. We observed significant enrichment of genes in the NOTCH pathway in the MZB-2 subset only (Fig. 4 C).

To understand further how these subsets relate to other key populations, we asked if either MZB subset might be associated with the age-associated ABC subset that expresses the transcription factor Tbet. Most notably the *Tbx21* gene that encodes Tbet was a feature of MZB-2 subset together with aNAV and DN-B in the spleen only (Fig. 4 D).

FcRL4 is an inhibitory receptor for aggregated IgA that is specifically induced on memory and MZB cells when in proximity to mucosal epithelium (Amara et al., 2017; Ehrhardt et al., 2005; Falini et al., 2003; Zhao et al., 2018). Here we see *FcRL4* gene expression by both subsets of MZB, aNAV, and ABC subsets 1 and 2 in the appendix. Interestingly, *FcRL4* gene was observed by only MZB-1 cells in the spleen (Fig. 4 E), consistent with the migration of this subset between spleen and the GALT microenvironment observed above (Fig. 3 C).

### MZB subsets in the MZ microenvironment

The term ‘marginal zone’ originally referred to the zone of B cells on the periphery of the white pulp and in the perifollicular region in human GALT (Mebius et al., 2004; Spencer et al., 1985). We therefore asked if the subset complexity in MZB we observe by analysis of cells in suspension could be observed in the splenic MZ.

We visualised microenvironments in splenic white pulp by imaging mass cytometry (Giesen et al.). The GC, composed of B cells, T cells, and macrophages, had areas of high B cell proliferation (Fig. 5 A). The GC was surrounded by a mantle zone of IgD^+^CD27^-^ naive B cells (Fig. 5 B). The mantle was in turn surrounded by IgD^+/-^CD1c^+^CD27^+^ cells, corresponding phenotypically to MZB cells identified in cell suspension (Fig. 5, B-D and Fig. S6 A). Visualisation of masked cells highlighted that the most peripheral B cell subset in the MZ was CD1c^-^, and likely corresponded to memory B cells, as also observed peripherally in GALT (Fig. 5, B and D). DDX21 expression, a feature of MZB-2 cell subset (Fig. 4 B), was significantly higher in CD1c^+^ than CD1c^-^ cells in the MZ (Fig. 5 E).

**Fig. 5:**
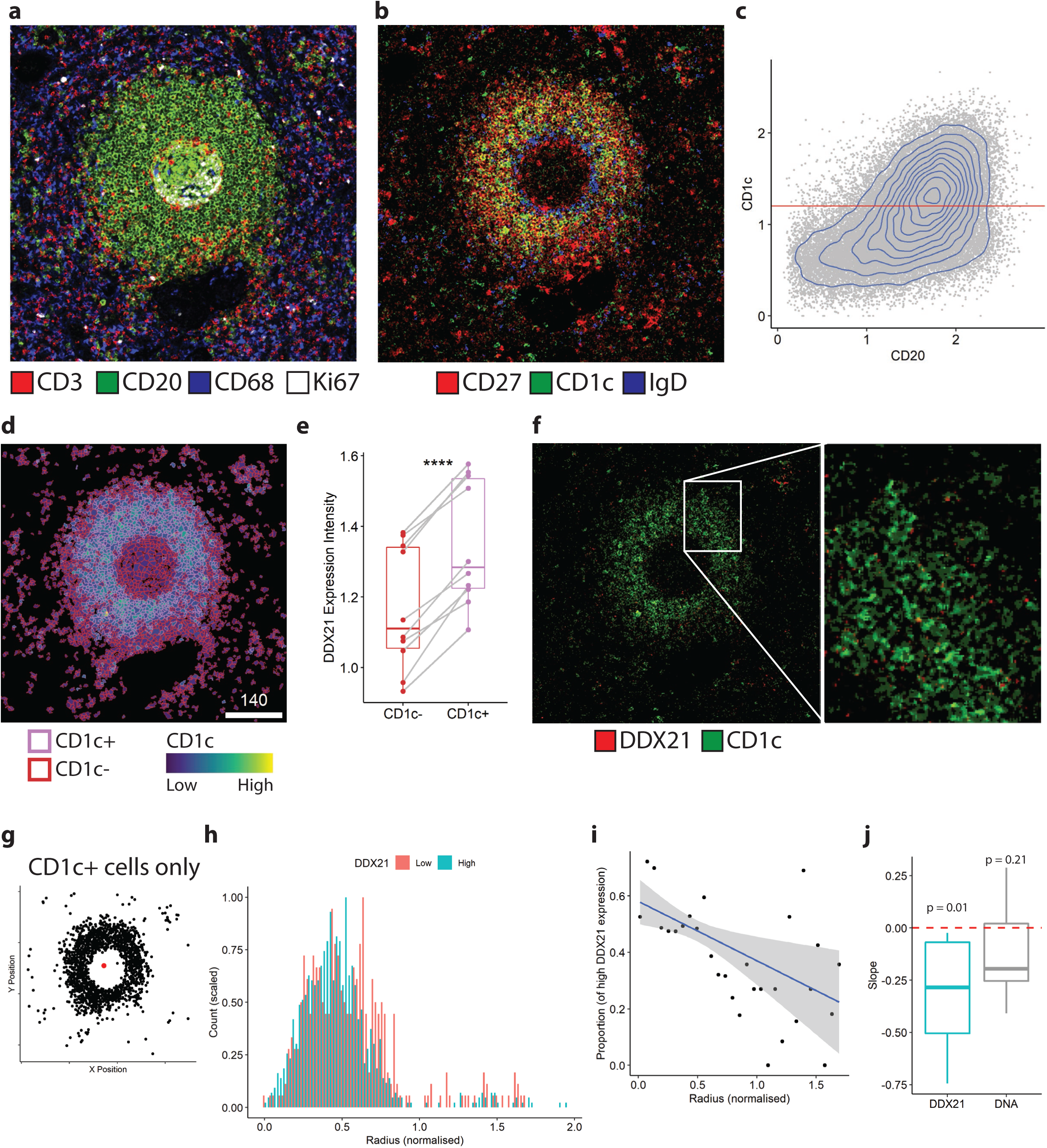
Spatial distribution of MZB in the spleen. Visualisation of microanatomy of human spleen by imaging mass cytometry. A representative example of spleen from 10 regions of interest. **(A).** Composite pixel-level image to visualise CD3 (T cells; red), CD20 (B cells; green), CD68 (macrophages; blue) and Ki67 (proliferation; white). **(B).** CD27 (red), CD1c (green), and IgD (blue). **(C).** CD1c gating threshold for segmented CD20^+^ single cells. **(D).** Visualisation of CD1c^+/-^ cell type (coloured by outline) and CD1c marker expression (coloured by fill) on CD20^+^ segmentation masks. **(E).** DDX21 expression levels between CD1c^-^ and CD1c^+^ B cells. Statistics for different expression levels between groups were assessed by paired t-test. **** p < 0.0001. **(F).** CD1c in green and DDX21 in red are visualised as a composite pixel-level image. **(G).** Scatterplot indicating the location of every CD1c^+^ B cell (black), and the centre point of these cells (red). **(H).** Distribution of the number of cells (y-axis; maximum value scaled to 1) at the various binned spatial distances between the centre point and every CD1c^+^ cell (x-axis; mean scaled to 0.5). Colour represents high (blue) and low (red) DDX21 expression. **(I).** Proportion of high DDX21 expression at each binned spatial distance from the previous histogram. A linear regression, weighted by cell counts, was performed (blue line). Grey indicates 95% confidence intervals. **(J).** Boxplot summary of the weighted linear regression slopes from regions of interest with a single follicle (n = 8) for DDX21 and DNA. One sample t-tests were used to compare the slopes for each marker to zero.

Consistent with expression of DDX21 by an MZB subset, we observed punctate nuclear and cytoplasmic DDX21 staining in a proportion of MZB cells (Fig. 5 F). We identified a distribution bias in DDX21 expressing CD1c^+^ cells in the MZ, with these being closer to the centre than cells lacking DDX21. Relative to the distance between a cell and the centre point (average X and Y position of all cells per image), the proportion of high and low DDX21 expression in CD1c+ cells was compared (Fig. 5, G-I and Fig. S6 B). If there was no spatial bias along the radial axis, the expected slope of the weighted linear regression would be zero. In other words, based on the linear regression slopes, DDX21 had a radial-biased distribution whereas DNA did not (Fig. 5 J).

In summary CD1c^+^ MZB and memory B cells occupy different locations within the MZ. In addition, of the intermingled CD1c expressing MZB subsets, DDX21^+^ MZB-2 cells locate significantly closer to the GC.

### MZB subsets in blood

We and others have previously studied MZB in blood and identified a differentiation pathway from IgM^hi^ TS cells that is reduced in severe systemic lupus erythematosus (SLE) (Tull et al., 2021). We therefore explored if either or both of the MZB subsets identified here are analogous to previously studied circulating MZB.

To address this, we analysed scRNAseq data acquired from sorted blood CD19^+^ B cells from 3 healthy controls and 3 patients with severe SLE (Fig. S7 A). Following normalisation and quality control, data from all 6 donors were pooled (Fig. S7, B-D), and then clustered according to 2000 variable genes. The 10 clusters identified according to gene and surface protein expression (Fig. S8, A and B) included 2 MZB clusters with the surface phenotype CD27^+^IgM^+^IgD^+^ and expression of *CD1C* and *PLD4* that are hallmarks of MZB cells (Fig S8C and Fig. 6 A-C). We named these clusters MZB-1 and MZB-2 by comparison with MZB-1 and MZB-2 identified in tissues above because of their relationships with other clusters and some transcriptomic features as described below.

**Fig. 6:**
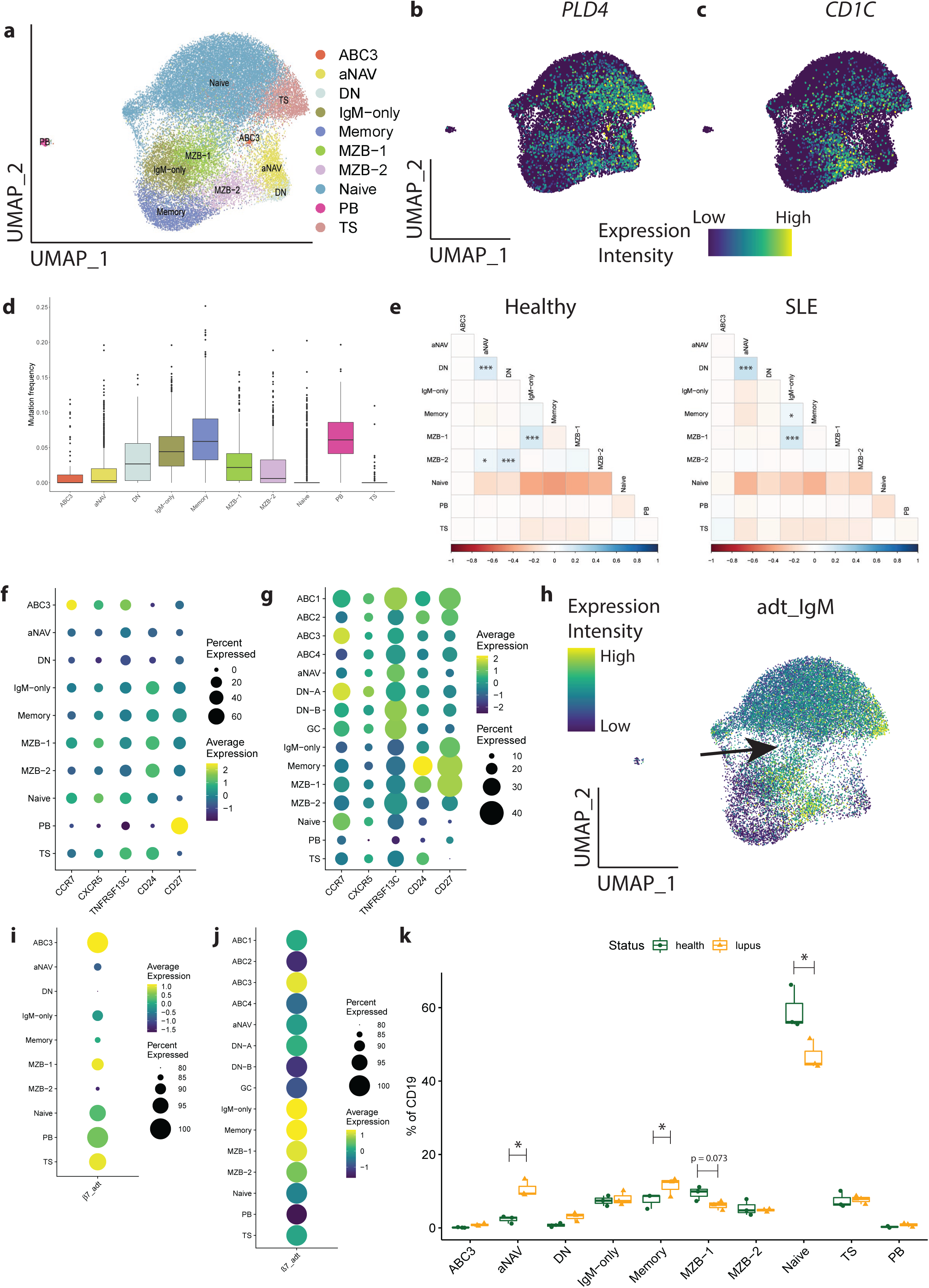
Comparison of MZB subsets in blood of healthy and SLE donors. **(A).** UMAP visualisation of B cell composition in blood coloured by B cell subsets, annotated based on Seurat unsupervised clusters (ABC3, activated B cells 3; aNAV, activated naive; DN, double negative; GC, germinal centre B cells; MZB, marginal zone B cells; PB, plasmablast; TS, transitional B cells). **(B).** UMAP visualisation of *PLD4* mRNA transcripts. **(C).** UMAP visualisation of *CD1C* mRNA transcripts. **(D).** Frequency of somatic mutations in *IGHV* associated with each B cell subset. **(E).** Clonal relatedness between B cell subsets within blood of healthy and SLE patients illustrated in a correlation plot. Significant relationships were assessed using Spearman correlation along with pairwise p-values that were then corrected using Holm’s method. **(F) and (G):** Dot plots illustrating the gene expressions of markers that discriminated between two subsets of B cells by mass cytometry in Fig. 1. **(F).** for B cell subtypes in healthy blood and **(G).** lymphoid tissues. **(H).** UMAP visualisation of IgM ADT surface protein expression in healthy blood B cells. Arrow indicates the IgM^hi^ bridge linking MZB-1 and naive B cells. **(I) and (J)**: Dot plots illustrating ADT surface protein expression of ß7 integrin for B cell subtypes in **(I).** healthy blood and **(J).** tissues. **(K).** Relative proportion of B cell subsets from healthy and SLE blood. Statistics for differential abundance between samples for each subset were assessed by estimated marginal means with Bonferroni correction for multiple comparisons. *p < 0.05, **p < 0.01, ***p < 0.001.

By analysis of the VDJ libraries, we identified that the frequency of SHM in each circulating subset was consistent with that observed in the corresponding subset in tissue (Fig. S9, A and B and Fig. 6 D). The MZB-1 subset had higher SHM than MZB-2 subset in blood.

In both health and severe SLE, there was a significant tendency for MZB-1 cells to have clonal relationships with IgM-only B cells (Fig. 6 E). In contrast, MZB-2 showed a significant tendency to be clonally related to aNAV and DN B cells in health. A clonal link between aNAV and DN was observed in SLE, consistent with published work(Jenks et al., 2018; Tull et al., 2021) (Fig. 6 E).

To cross reference data acquired by mass cytometry and single cell transcriptomic analyses of tissues and blood, we compared gene expression for those proteins that distinguished the two subsets of tissue MZB in the original mass cytometry analysis (Fig. 1 C); this gene set also discriminated MZB-1 and MZB-2 in tissues and blood (Fig. 6, F and G).

We have recently described MZB derived from IgM^hi^ TS B cells that is associated with gut homing and that is depleted in severe lupus (Tull et al., 2021). We therefore asked where this pathway sits in the context of MZB-1 and MZB-2. Whilst both subsets of MZB in blood were IgM^hi^, the MZB-1 subset was visibly linked to the naive B cell pool by an IgM^hi^ bridge as described previously (Fig. 6 H). In addition, expression of gut-homing ß7 integrin was associated with MZB-1 in both blood and tissues (Fig. 6, I and J).

We next compared the frequencies of cells in subsets in health and SLE. We observed that MZB-1 cells tend to be more abundant than MZB-2 cells in the blood in health, but that the frequency of the MZB-1 subset was consistently reduced in severe SLE (Fig. 6 K). As in previous studies, DN, aNAV and class switched memory B cells were increased in frequency in SLE(Jenks et al., 2018; Tull et al., 2021) (Fig. 6 K).

## Discussion

Here we describe subsets of human B cells in matched appendix, mLN and spleen samples from deceased transplant donors using multiparameter unsupervised methods. In addition to expected and novel tissue-associated subsets of activated B cells, we observed two subsets of MZB, both of which are at highest frequency in the spleen, where they are observed in the MZ. Of these, the MZB-1 subset was the most highly migratory subset overall in terms of both spread of clones between tissues and in relative frequency in blood. This subset was clonally related to IgM-only B cells that were also highly mobile.

Our recent analysis of circulating human B cell subsets identified IgM^hi^ intermediary stages in the development of human MZB cells from TS B cells (Tull et al., 2021). Here we refine those findings, by showing that the IgM^hi^ pathway is only relevant to the MZB-1 subset. In comparison, the MZB-2 subset has a signature of NOTCH related genes, suggesting that this subset matures by NOTCH ligation, as described previously in humans and mice (Descatoire et al., 2014; Saito et al., 2003). There was no evidence in this study that the two MZB subsets are developmentally related since they are largely clonally separate, with the MZB-2 subset having a more diverse repertoire, less V region mutations, and with its clones tending to localise to single tissue sites.

The MZ is traditionally associated with innate like immune responses to bacterial antigens with repeating subunit structures: so-called T-independent type 2 antigens. Children less than 2 years of age (when MZB cells are poorly developed and consist of non-clonal cells with low mutation loads) and also individuals who were splenectomised early in life are particularly at risk of infections with bacterial pathogens (Amlot and Hayes, 1985; Timens et al., 1989; Weller et al., 2008). Patients with severe SLE are also susceptible to infection with pneumococci, suggesting that it is principally the MZB-1 population (which is depleted in severe SLE) that provides protection (Danza and Ruiz-Irastorza, 2013).

In contrast, the MZB-2 subset, expressed the RNA helicase DDX21 and may have an anti-viral role (Chen et al., 2014). The MZB-2 subset when in spleen was enriched in cells expressing transcription factor Tbet; these have been shown to have anti-viral properties in mice (Johnson et al., 2020). As in mice, the splenic Tbet-expressing subset had a tendency to remain local. Low levels of mutations also make them similar to mouse MZB where MZB heterogeneity has also been previously observed (Dammers et al., 2000; Song and Cerny, 2003). When members of this subset were detected in blood, however, they did show clonal relationships with aNAV and DN cells that have been linked to the extrafollicular response, suggesting that the cells that cluster together may have different functional phases when static or migratory, that may be dependent on local interactions such as NOTCH2 ligation (Tipton et al., 2015; Weller et al., 2004).

Whereas the importance of IgA in mucosal immunology is well defined, the role of IgM is less clear. There are both memory and clonally related plasma cells that express only IgM in human gut, suggesting that IgM-only cells are not merely recently activated naive B cells (Magri et al., 2017). The high number of mutations in the IgM-only B cell subset supports this concept. We have previously identified that CD27^+^IgM^+^IgD^+^ cells in GALT can be clonally related to IgM-only cells (Zhao et al., 2018). However, IgM-only cells can be clonally related to class switched memory cells whereas CD27^+^IgM^+^IgD^+^ cells rarely are (Zhao et al., 2018). This suggests that IgM-only cells in tissues are heterogenous as a population. Here we identify IgM-only cells as a distinct transcriptome driven cluster, with MZB-1 relatives and clone members that are disseminated between the three tissues. Of note, many IgM-only cells were clustered alongside IgA and IgG class switched memory cells, consistent with distinct functional groups classified as IgM-only cells that we do not yet understand.

Variants of blood CD27^+^IgM^+^IgD^+^ B cells defined according to levels of IgM expression have been identified (Bautista et al., 2020). Differences in IgM expression was not a feature of the subsets identified in tissues in this study, and we saw no evidence of enrichment of genes that discriminated between the subsets described.

The GC cells, which were most abundant in appendix had some surprising features, including a tendency of clonal relatives to remain local. Also surprising was the dominant use of the IgM isotype and the relative abundance of cells with low mutation frequencies. These features are at odds with the reputation of GALT GCs as the hub that constantly replenishes IgA, and suggests that the role of GALT GCs in maturation of the naive repertoire and in sustaining IgM only responses may have been underestimated (Roco et al., 2019).

We observed small clones of TS and naive B cells in appendix and spleen. These could only be observed within single sites of lymphoid tissue suggesting that when they join the systemic circulating pool, clone members become separated and no longer detectable due to dilution. When light chains were captured in addition to the heavy chains by which clones were identified, these were identical between clone members supporting the concept of local proliferation of cells that were already fully mature in terms of BCR rearrangement. Activated TS cells have been observed previously in GALT (Vossenkämper et al., 2013). Selection occurs as B cells mature from TS to naive follicular and MZ cells, and it is possible that checkpoints in appendix and spleen involving cell division are involved in shaping the naive B cell repertoire (Wardemann et al., 2003).

We identified 4 subsets of ABC that are likely to represent different activation states or stages of differentiation. Only one, ABC3, was also present in blood. ABC1 is genotypically like GC B cells; however, it does not express classic GC markers such as *BCL6*. In addition, while clonally related to GC cells, the ABC1 subset have higher mutation frequency, suggesting that it is a more terminally differentiated subset. Thus, the ABC1 subset could reflect cells that have just exited the GC, possibly on their way to undergo class switch recombination, but have not undergone further class-switching.

It is possible that the two subsets of MZB that we describe are benign analogues of subtypes of MZB lymphomas. MZB-1 may represent benign analogues of marginal zone B cell lymphoma of mucosa-associated lymphoid tissue (MALT lymphoma), a malignancy that can be driven by bacterial infections (Wotherspoon et al., 1993). These tumours tend to express IgM and can circulate from GALT via blood to the spleen where they tend to occupy the MZ (Spencer et al., 1990). Like MZB-1, MALT lymphomas can express FcRL4 when encountered in the mucosal microenvironment (Falini et al., 2003). Here we show that FcRL4 transcripts can be retained by MZB-1 in spleen consistent with migration between gut and spleen. In contrast MZB-2 may represent the benign analogues of splenic marginal zone lymphomas (SMZL). Cases of SMZL tend to localise to the spleen but not other secondary lymphoid tissues; cellular variants may express the transcription factor Tbet(Lohneis et al., 2014); and they may arise through translocations involving the NOTCH pathway, activation of which is observed here in MZB-2 only (Rossi et al., 2012). Understanding benign analogues of lymphoma subtypes may help future identification of drivers and thus potential therapeutic pathway inhibitors.

Overall, the deep analysis of B cells in tissues that we present, which combined undirected methods of grouping of similar cells with knowledge and reference-based subset alignments to the groups identified, provides a more accurate vision of tissue-based subsets and their interrelatedness within and between tissues than was previously available. There were organ-specific expression patterns within subsets, demonstrating that the local microarchitecture and milieu will determine cellular functions. MZB cells have been referred to as the ‘elephant in the room’ of human B cell biology (Nemazee, 2021). The clarity that we provide here dismisses much of the mystery and confusion in this field. The resolution of existing data solves the many controversies within the field and provides a reference platform going forwards.

## Methods

### Experimental subject details

Human tissue was obtained from deceased adult transplant organ donors with research ethics committee (REC) approval and informed consent from the donor family (reference 15/EE/0152, East of England Cambridge South Research Ethics Committee). Studies of human tissues were approved by London, Camberwell St Giles Research Ethics Committee (study 11/LO/1274 Immunology of the intestine; features associated with autoimmunity).

SLE patients were recruited using the following criteria: (1) fulfilment of four or more revised American College of Rheumatology classification criteria, (2) ANA-positive, (3) biological (belimumab or rituximab) naive, and (4) immunosuppressive regimen does not include azathioprine or cyclophosphamide within 6 months of sample collection due to the severe depletion of naive B cells by these medications. All LN patients had diagnostic confirmation by renal biopsy. Blood was obtained from SLE patients and healthy controls with informed consent and REC approval (REC reference 11/LO/1433: Immune regulation in autoimmune rheumatic disease, London–City Road & Hampstead Research Ethics Committee).

### Sample processing

Matched appendix, mLN, spleen were from deceased transplant organ donors and stored in UW (University of Wisconsin) solution at 4°C. All tissue preparation and lymphocyte isolation procedures were performed with RPMI-1640 containing heat inactivated 10% FCS, 2 mM L-glutamine, 100 IU/mL penicillin and 100 μg/mL streptomycin (RPMI-P/S) unless stated otherwise. Appendixes (cut into 1-2mm pieces) were incubated at 37°C for 30 minutes in medium with 1 mg/mL collagenase IV (Sigma-Aldrich) and 1 mg/mL DNAse I (Roche). Lightly digested appendix as well as fresh mLN were then teased through a 70 μm nylon cell strainer, washed with medium, and re-suspended for cryopreservation. Small sections of spleen were placed in gentleMACS C tubes (Miltenyi) topped up with PBS + 2% FCS where the gentleMACS setting B ran three times in the gentleMACS Dissociator. The solution was then filtered through a 70 μm nylon cell strainer (diluted with PBS as necessary). Spleen solution was then layered onto Lymphoprep (Stemcell Technologies) according to manufacturer’s instructions and then centrifuged at 800 x g for 20 minutes at 4°C with no brakes. The buffy coat was then collected, washed with medium, centrifuged, and re-suspended in red blood cell lysis buffer (0.17M ammonium chloride) for 7 minutes at room temperature to lyse erythrocytes. This suspension was topped up to 20 mL, centrifuged, and re-suspended in freezing medium. Tissue cells were cryopreserved in a freezing medium of FCS + 10% dimethyl sulfoxide (DMSO) at a concentration of 1×10^7^ whenever possible.

Blood samples were diluted 1:1 in RPMI-P/S, and then layered onto Ficoll and centrifuged for 25 minutes with no brake. The buffy coat layer was collected, and cells were washed. Peripheral blood mononuclear cells (PBMCs) were cryopreserved in FCS + 10% DMSO at a concentration of 1 x 10^7^ whenever possible.

### Mass cytometry

Frozen samples were thawed in a 37°C water bath and washed in 2 mL of RPMI-1640 + 10% FBS, 0.1 mg/mL DNase I (Roche), and transferred to a larger volume of 10 mL. After centrifuging at 400 x g for 7 minutes at 37°C, samples were resuspended in 2 mL of media as above and rested for 30 minutes at 37°C.

Cell staining medium (CSM) consists of low barium PBS with 2.5 g bovine serum albumin (BSA; Sigma) and 2 mM ethylenediaminetetraacetic acid (EDTA; Invitrogen). Perm buffer (CSM-S) includes CSM (as above) with 1.5 g saponin. For live/dead staining, approximately 4 million cells were used after resting. Samples were topped up with 10 mL of pre-warmed media (as above) and centrifuged at 400 x g for 7 minutes at 37°C. Cells were then resuspended in 1 mL of diluted rhodium-intercalater (stock diluted 1:500 in PBS) and incubated at room temperature for 20 minutes. Samples were then topped up with 10 mL of CSM, centrifuged at 400 x g for 7 minutes at 37°C, and counted. For antibody staining, 2 million cells were used. Samples were resuspended in 40 μL CSM with 5 μL Fc block, and incubated on ice for 10 minutes. Volumes were then adjusted to 99 μL CSM, 1 μL of IgG was added to each sample and incubated on ice for 30 minutes. After washing with CSM and centrifuged, samples were resuspended in the mastermix as described in Supplementary Table 1 in a total volume of 100 μL and incubated on ice for 30 minutes. Samples were then washed, centrifuged and resuspended in 1 mL of 2% paraformaldehyde in PBS overnight at 4°C. The following day, cells were washed in 1 x PBS and DNA was stained with 1 μM Intercalatin in 500 μl permeabilization buffer at room temperature for 20 minutes. Cells were then washed twice in 1 x PBS, twice in Milli-Q water before being resuspended in Milli-Q water plus EQ beads to a concentration of 0.5 x 10^6^ /ml and run on a Helios mass cytometer.

### Pre-processing and normalisation of mass cytometry data

FCS files were normalised using bead standards and the Normalizer program developed by Nolan’s group (v0.3, available online at https://github.com/nolanlab/beadnormalization/releases) (Finck et al., 2013). The result of the bead normalization is visualized in Fig. S1 A. The mass cytometry data was initially manually gated in Cytobank (https://mrc.cytobank.org/) using the gating strategy shown in Fig. S1 D to identify live CD19^+^ B cells for downstream analysis.

For batch normalisation purposes, each paired tissue set was run in the same batch with an internal biological sample from the same spleen as reference. Internal controls between batches were normalised by transforming the pooled intensity distribution of all batches towards a reference distribution (Lun et al.). The range normalisation method scaled the marker intensities so that the distribution range was the same for each batch. By comparing and matching the 1st and 99th percentiles of each batch to the reference distribution, a linear function was then defined and applied to all markers of all samples during normalisation. The effects of normalisation on the internal controls and the entire dataset of select markers is visualised in Fig. S1 E and F.

### Differential abundance analysis of mass cytometry data

After the data were normalised, cells were allocated into hyperspheres, and then tested for differential abundance between tissues for each hypersphere while controlling for false discovery rate (Lun et al., 2017). Hyperspheres represent cells in regions of the data with similar marker phenotype. Importantly, unlike clustering, a cell can be allocated into more than one hypersphere. Low-abundance hyperspheres with average counts below 100 were filtered out (Fig. S1 G). Multidimensional scaling plot was used to determine if abundance differences were attributed to different tissue type and not biological variation (Fig. S1 H). As developed previously by Lun *et al*., empirical Bayes method was used to allow hypersphere-specific variation estimates even when replicate numbers were small by borrowing distribution information between hyperspheres (Robinson et al., 2010).

To compare the cell counts between samples, significant differences in cell abundance between conditions were tested. The null hypothesis of no change in the average counts of cells between conditions for each hyperspheres was tested using negative binomial generalised linear model (NB GLMs) implementation in edgeR package (Robinson et al., 2010). Finally, to control false discovery rates, spatial false discovery rate of 5% was used; this is the volume of differentially abundant hyperspheres that are false positives and considers the density of hyperspheres.

### Cell sorting and CITE-seq antibody staining

Cryopreserved samples were thawed in a 37°C water bath and washed in RPMI with penicillin and streptomycin. After cell counting, 3 million cells per sample re-suspended in PBS with 1% BSA were transferred to Eppendorf LoBind Microcentrifuge tubes and washed. Cells were then incubated for 10 minutes at 4°C with True stain Fc block (BioLegend) and washed. Cells were stained on ice for 30 minutes with a mastermix consisting of pre-titrated concentrations of flow cytometry antibody (CD19 PerCP-Cy5.5, 1:50) as well as CITE-seq antibodies at a concentration of 8 μg/mL shown in Fig. S2 B. Cells were then washed two times before cell sorting. Viability staining with DAPI 0.1mg/ml diluted 1:1000 was added prior to sample acquisition on the flow cytometer. Anti-Mouse/Rat beads (BD Biosciences) were used for fluorescent compensation and gates were set using appropriate isotype controls. Cell sorting was performed using a BD FACSAria (BD Biosciences) where live single CD19^+^ B cells were sorted (Fig. S2 A). Samples were then washed three times before loading onto the 10X Chromium controller.

### Imaging mass cytometry

A panel containing 10 metal-tagged antibodies (Supplementary Table 2) was designed to identify and characterize immune populations expected to be present in the splenic white pulp. Each antibody was tested in spleen section at different concentrations and the resulting IMC images were visually examined to select the best concentration. Formalin Fixed Paraffin Embedded (FFFPE) samples of human spleen were obtained from three anonymous healthy adult donors, cut in 4 μm-thick sections using a microtome and subjected to IMC staining. Briefly, sections were deparaffinized, rehydrated and subjected to antigen retrieval using a pressure cooker. Tissues were then blocked and incubated with the mix of metal-conjugated antibodies contained in the panel overnight at 4°C and subsequently incubated with the DNA intercalator Iridium Cell-ID™ Intercalator-Ir (Fluidigm) before being air-dried. Slides were then inserted into the Hyperion Imaging System and photographed to aid region selection. A total of 10 regions of approximatively 1mm^2^ were selected to contain at least one identifiable GC and subsequently laser-ablated at 200 Hz frequency at 1μm/pixel resolution.

### Imaging mass cytometry analysis

Single cell segmentation was performed prior to single cell protein expression analysis. Using CellProfiler, staining for DNA was used to identify cell nuclei and B cells were detected by masking nuclei which also contained CD20 (Mcquin et al., 2018). Resulting images showed morphologically appropriate cell boundaries and centres. Cell locations and measurements of staining intensity for the various markers within each cell were recorded. Pixel-level composite images were created using histoCAT (Schapiro et al., 2017). Cytomapper was used to overlay the single cell metadata onto the cell segmentation masks (Eling et al., 2020). Marker expressions were scaled by arcsinh with a factor of 0.1 for visualisation purposes. The CD1c threshold of 1.2 was used. The centre point for each image was determined using the average X and Y position of every CD1c^+^ cell. Images with multiple follicles were excluded from downstream analysis. The distance between the centre point and each CD1c^+^ cell was calculated. This distance distribution was scaled such that the overall radius of each image was 1. In the histogram, counts were scaled by image such that the maximum count was 1 and the bins for the spatial distances were 0.05. Marker expression for each cell was classified as high or low if they were above or below the median marker expression per image. Linear regression was weighted by the cell counts in each bin. Paired and one sample t-tests were performed and illustrated using ggpubr (Kassambara, 2020a).

### Single cell RNA sequencing library preparation

Sorted CD19^+^ cell populations were loaded onto a 10X Genomics Chromium Controller and the libraries (5’ gene expression, VDJ, ADT) were prepared according to manufacturer’s guidelines. The Illumina HiSeq 2500 High Output platform was used to sequence the samples. Transcript alignment and generation of feature-barcode matrices for downstream analysis were performed using the 10X Genomics Cell Ranger workflow (Fig. S2 C).

### Single cell transcriptome analysis of tissues

Using the Seurat R package (Version 3.2.2), sorted CD19^+^ cells with high mitochondrial transcripts, low/high number of unique genes per cell, and low/high total RNA transcripts were filtered out (Stuart et al., 2019). The threshold used was 3 times the mean absolute deviation of each sample (Fig. S2, D-F). B cells were isolated based on the expression of B cell specific genes (*CD79A, CD79B, CD19, MS4A1*) and absence of T cell specific genes (*CD2, CD3D, CD3E, CD3G, CD4, CD8B, CD7*), and *TCR* genes. Genes that were only expressed in 10 cells or fewer in the entire dataset were filtered out. In addition, *IGHV* and *TCR* genes were removed prior to downstream analysis. The data was transformed and integrated using the SCTransform and Integration workflow (Fig. S2 G). For more efficient clustering and dimension reduction, PCA was performed on the top 3000 transcriptomic variable features, and the top 25 principal components were used for downstream analysis. Cells were clustered using the standard Seurat graph-based clustering approach with resolution of 1.3. Genes were deemed significantly differentially expressed using the Wilcoxon rank sum test with genes that have a log fold change threshold of 0.25 and expressed in 10% of either population (FindClusters function within Seurat). Clusters with similar differentially expressed gene lists were combined, and all the clusters were annotated using the gene lists shown in Fig. S3.

A two-way ANOVA was conducted to examine the effects of subsets and tissue type on cell proportion. Residual analysis was performed to test for the assumptions of the ANOVA. Outliers were visually assessed using box plots, normality was assessed using Shapiro-Wilk’s normality test and homogeneity of variances was assessed by Levene’s test. There were no extreme outliers (points 3 times the interquartile range), residuals were normally distributed (p > 0.05) and there was homogeneity of variances (p > 0.05). Consequently, an analysis of simple main effects for subsets was performed with statistical significance receiving a Bonferroni adjustment. All pairwise comparisons (estimated marginal means) were analyzed between the different tissue types organized by subsets, and p-values were adjusted using Bonferroni (Kassambara, 2020b).

Gene set enrichment analysis was done using the Broad Institute’s GSEA software(Mootha et al., 2003; Subramanian et al., 2005). All relevant Notch gene sets were downloaded from the Molecular Signatures Database (MSigDB) and corrected for our dataset’s background. Spliced and unspliced RNA count matrices were obtained by scVelo (La Manno et al., 2018). The proportions of spliced and unspliced sequences were the following: appendix (33% spliced, 67% unspliced), mLN (31% spliced, 69% unspliced), and spleen (33% spliced, 67% unspliced). RNA velocity was calculated using scVelo(Bergen et al.).

### Single cell transcriptome analysis of blood

A similar workflow as described above was performed on sorted CD19^+^ blood from healthy and SLE patients with the following modifications. Genes that were only expressed in 3 cells or fewer in the entire dataset were filtered out. The standard Seurat integration workflow was used to normalise and integrated the samples. For more efficient clustering and dimension reduction, PCA was performed on the top 2000 variable features, and the top 20 principal components were used for downstream analysis. Cells were clustered using the standard Seurat graph-based clustering approach with resolution of 1.8.

### Single cell BCR analysis

The BCR repetoire analysis was done using the Immcantation framework (Version 4.0) (Gupta et al., 2015). V, D, and J genes were assigned using IgBLAST. Non-productive sequences are removed. Clonal threshold at 0.15 or 0.1 was determined from the Hamming distance (Fig. S4 B and 8 B). To normalize the donor effect when looking at the ratio of disseminated and resident clones, the ratio in each subset was divided by the average dissemination ratio in that donor. To determine the clonal correlation within and between tissues, the Spearman correlations per donor along with the pairwise p-values were calculated. These p-values were then corrected for multiple interference using Holm’s method in the R package RcmdrMisc (Fox, 2020).

### Data analysis and visualisation

All data analysis and visualisation were done in R (3.5.2, 4.0.2) unless stated otherwise (R Core Team, 2020). Dimension reduction with Uniform Manifold Approximation and Projection (UMAP) was implemented using uwot package (Melville, 2020). Heatmaps were visualised using ComplexHeatmap package (Gu et al., 2016). Circos plots were built using circlize package (Gu et al., 2014). Volcano plot was drawn using EnhancedVolcano package(Blighe et al., 2020). Correlation plots were drawn using corrplot package (Wei and Simko, 2017). Other single cell sequencing figures were drawn using Seurat (3.2.2), ggplot2, and ggpubr packages (Kassambara, 2020a; Stuart et al., 2019; Wickham, 2009).

### Data and code availability

All raw and processed next-generation sequencing data have been deposited with GEO under accession numbers XXXXX and XXXXX. Mass cytometry and image mass cytometry data available upon request. Code is available on Github (https://github.com/jspencer-lab/Siu_MZManuscript_2021).

## Acknowledgments

We thank the deceased organ donors, donor families and the Cambridge Biorepository for Translational Medicine for access to tissue samples. This work was funded by the Medical Research Council of Great Britain (MR/R000964/1 and MR/P021964/1), the Lupus Trust, the Wellcome Trust (220872/Z/20/Z), and the Chan Zuckerberg Initiative. We acknowledge support from the Flow Cytometry and Genomics Research Platforms within the Biomedical Research Centre at Guy’s and St Thomas’ NHS Trust and the King’s Health Partners and Cancer Research UK KHP Cancer Centre. We would also like to thank Silvia Cellone Trevelin for help with the histology, and Francesca Ciccarelli and Michele Bortolomeazzi for guidance with image analysis. J.H.S is supported by the Gates Cambridge Trust and the Canadian Centennial Scholarship Fund. M.B. is supported by research funds from The Swedish Research Council and the County Council of Västra Götaland.

## Author Contributions

Conceptualization and design of study: J.H.S., J.S., G.J.P, T.J.T; Sample identification and collection: J.H.S., K.T.M., K.S-P., G.J.P.; Data acquisition and methodology; J.H.S., T.J.T., L.M., C.W.A., R.E., P.D., U.D.K., M.K., K.T, J.S.; Data analysis: J.H.S., M.J.P., W.G., T.J.T; Supervision and funding: J.S, G.J.P., M.B, D.D’C. Writing the manuscript; J.H.S., J.S., G.J.P, M.B, T.J.T.

---

**Fig. S1:**
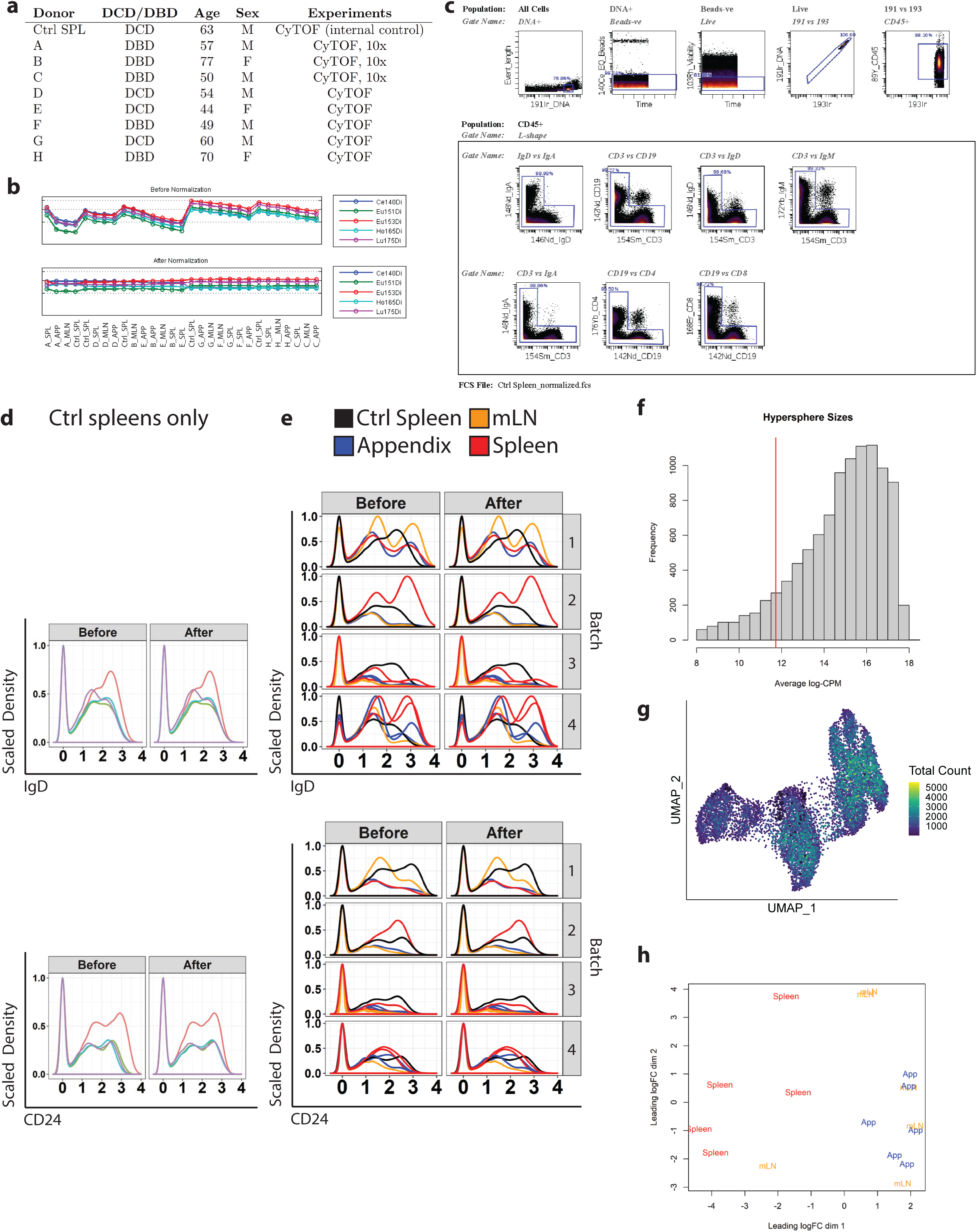
Donor details, quality control, preliminary gating, batch normalisation, and preliminary analysis of hyperspheres for mass cytometry. **(A).** Donor id, type of donation (donation after circulatory death, DCD; donation after brainstem death, DBD), age, sex, and experiments used. **(B).** Bead normalisation. **(C).** Preliminary gating for CD19^+^ B cells. Subsequent doublet gates (L-gates) excluded remaining minority of cells with implausible immunoglobulin and B/T cell combinations. **(D).** Density plots showcase the effects of range normalisation before and after on the same internal spleen control. Colours represent the same spleen in different batch runs. Range normalisation scaled marker intensities such that the distribution range was identical between batches. **(E).** Density plots showcase the effects of range normalisation before and after on the entire dataset. Colours represent different tissues: control spleen (black), appendix (blue), mLN (orange), spleen (red). **(F).** Frequency plot of hypersphere sizes. Red line indicates cut-off value for downstream analysis. **(G).** UMAP coloured by total cell counts for the three tissues (median total value of the donors) within each hypersphere. **(H).** Multidimensional scaling (MDS) plot of raw hypersphere sizes, coloured by tissue.

**Fig. S2:**
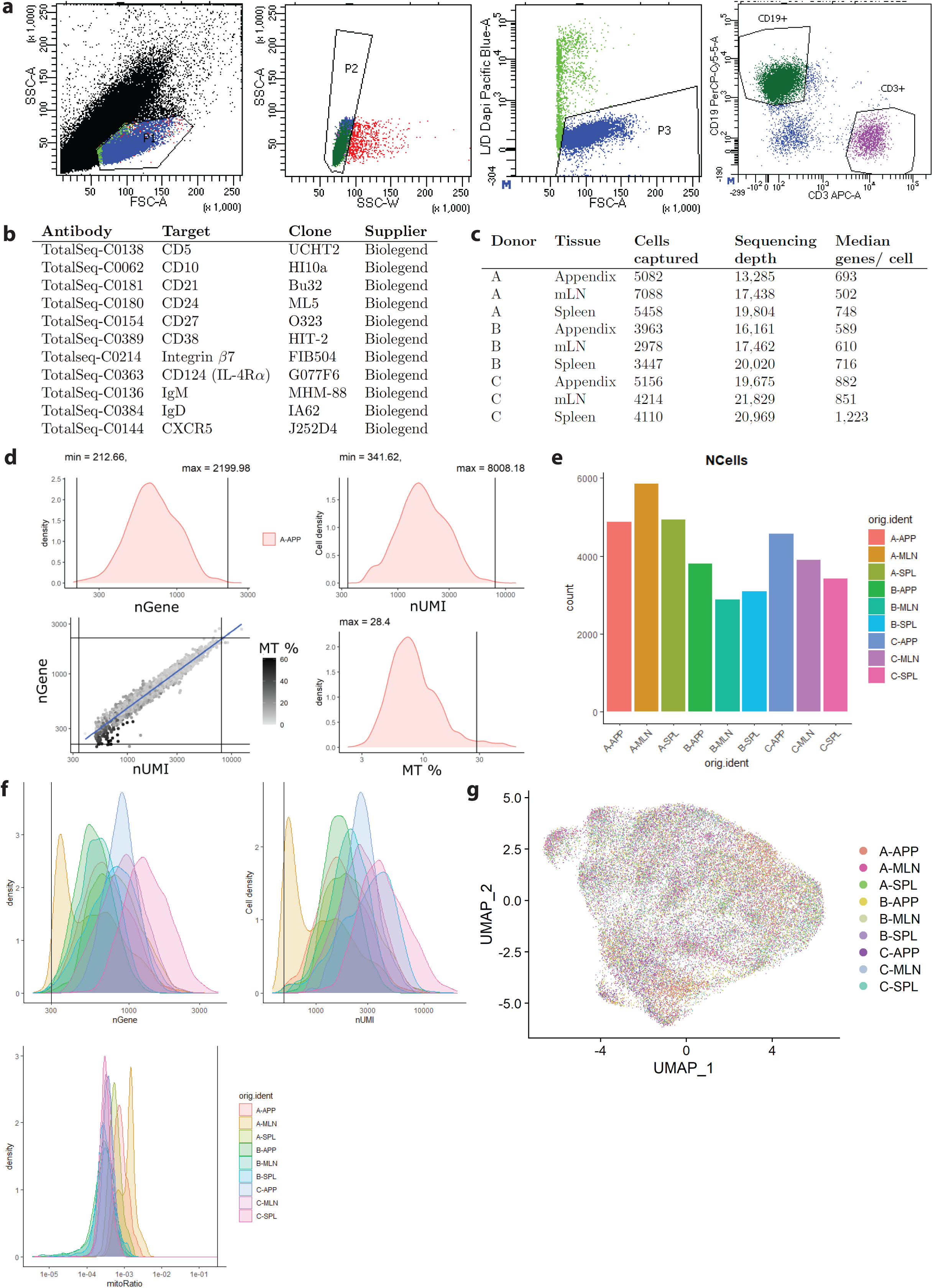
Sort strategy, 10X genomics workflow & quality control, and batch integration for single cell transcriptomics. **(A).** Gating strategy to sort live CD19^+^ B cells. **(B).** Total-Seq antibodies and clones used for surface labelling of CD19^+^ B cells. **(C).** Cells captured and sequencing depth of the three donors and their matched tissues. **(D).** Cell barcodes processed where barcodes with transcripts counts per cell (nUMI, unique molecular identifiers) and genes detected per cell (nGene) values above/below 3 median absolute deviation (MAD) and mitochondria percentage above MAD were removed. **(E).** Cell counts after quality control processing. **(F).** Summary of samples after auto processing. **(G).** UMAP visualisation of B cell composition in tissues after integration, coloured by sample.

**Fig. S3:**
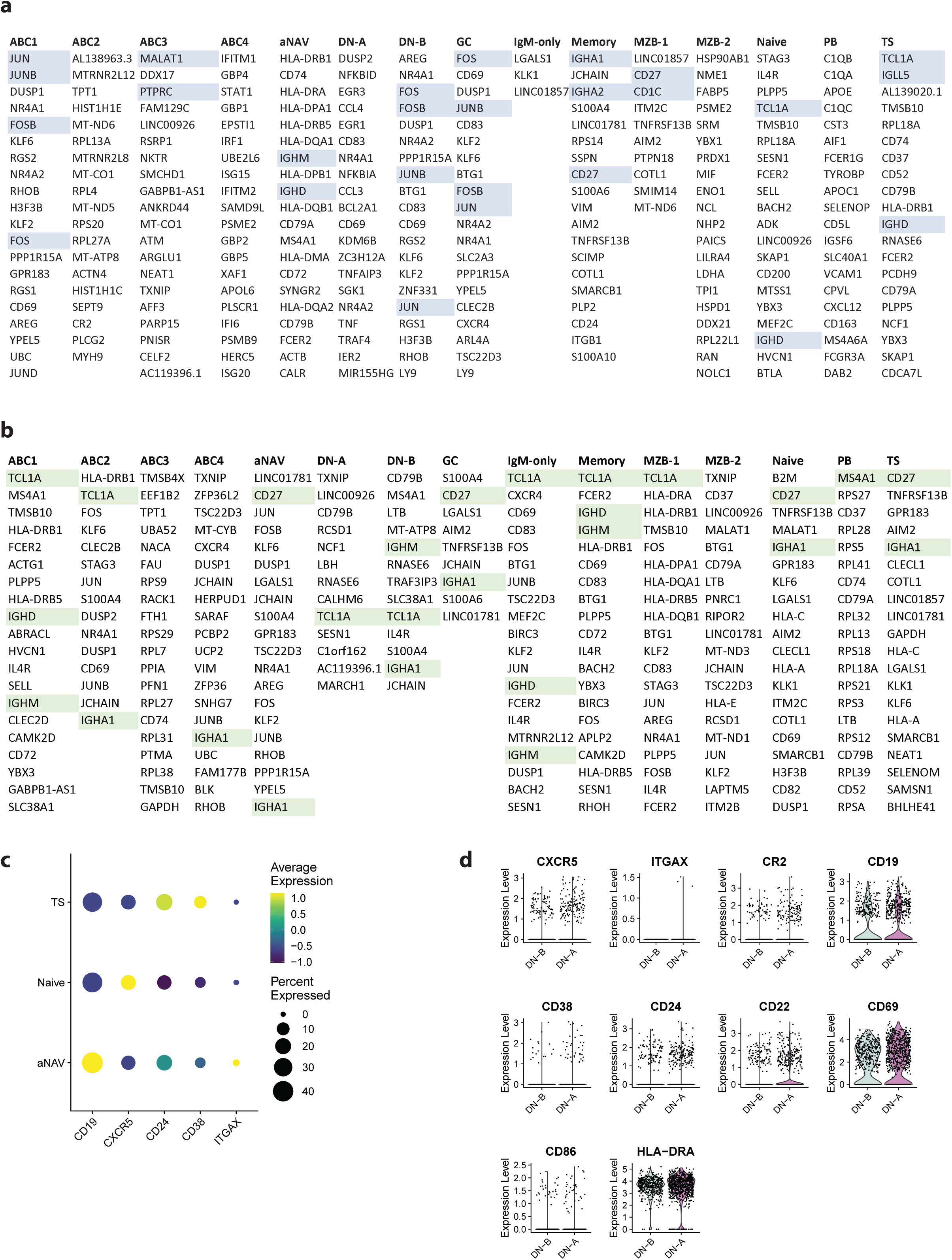
Differentially expressed gene for each B cell subtype in tissues. **(A).** Top 20 positive differentially expressed genes for each subtype. **(B).** Top 20 negative differentially expressed genes for each subtype. Genes used for annotating cluster are highlighted in blue and green. P values were calculated using Wilcoxon test with Bonferroni correction for multiple comparisons. **(C).** Dot plot illustrating *CD19, CXCR5, CD24, CD38, ITGAX (CD11c)* gene expression for TS, naive, and aNAV B cell subsets. **(D).** Violin plots of key DN differentiating genes for DN-A and DN-B subtypes.

**Fig. S4:**
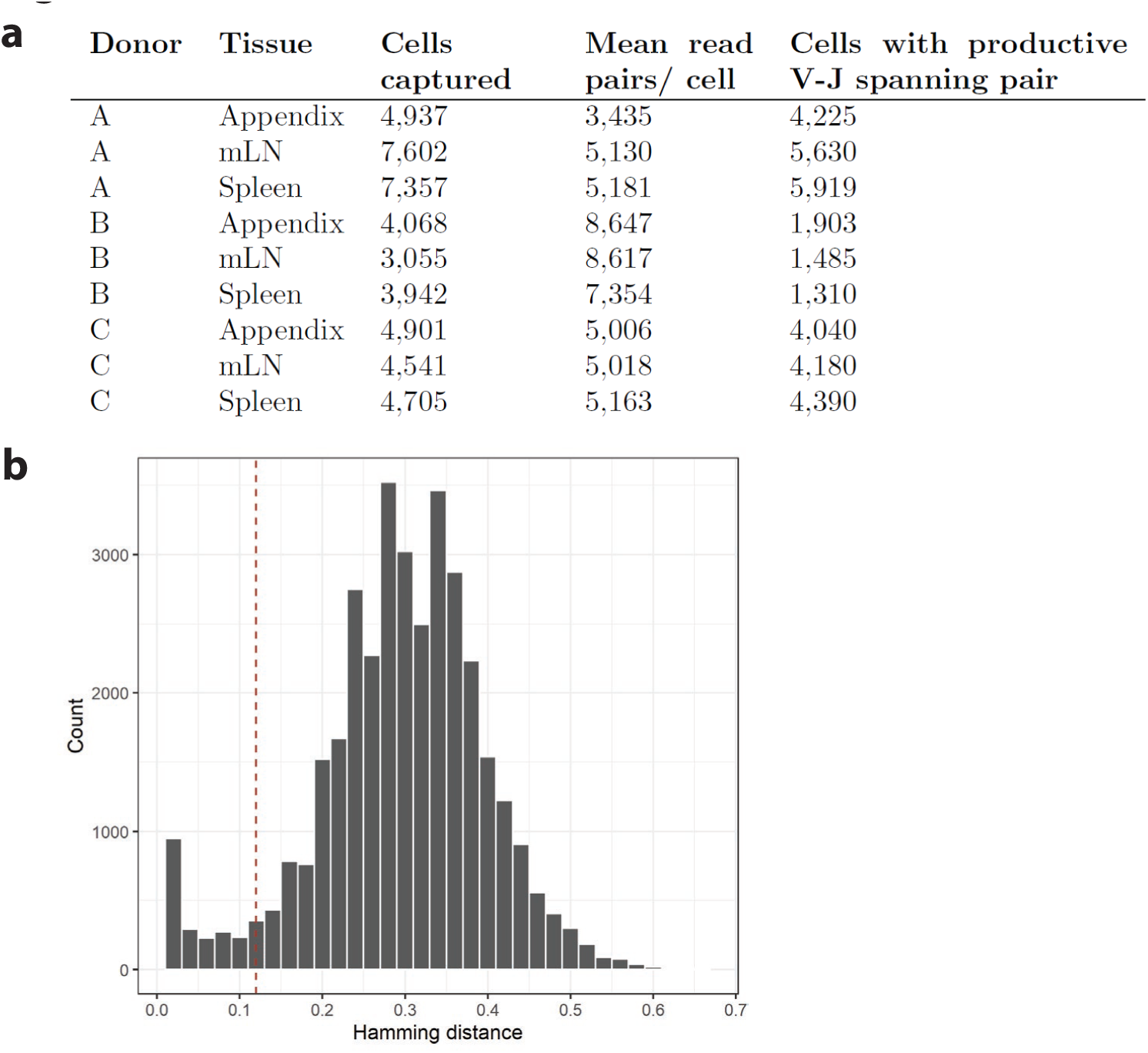
BCR workflow and clonal clustering threshold of tissues. **(A).** Cells captured, mean read pairs per cell, number of productive V-J cells for each sample. **(B).** Using the Immcantation pipeline, the distance between sequences and its nearest-neighbour was calculated to determine the threshold for separating clonal groups. The threshold (red dashed line) of 0.15 was determined by inspection of the distance-to-nearest plot.

**Fig. S5:**
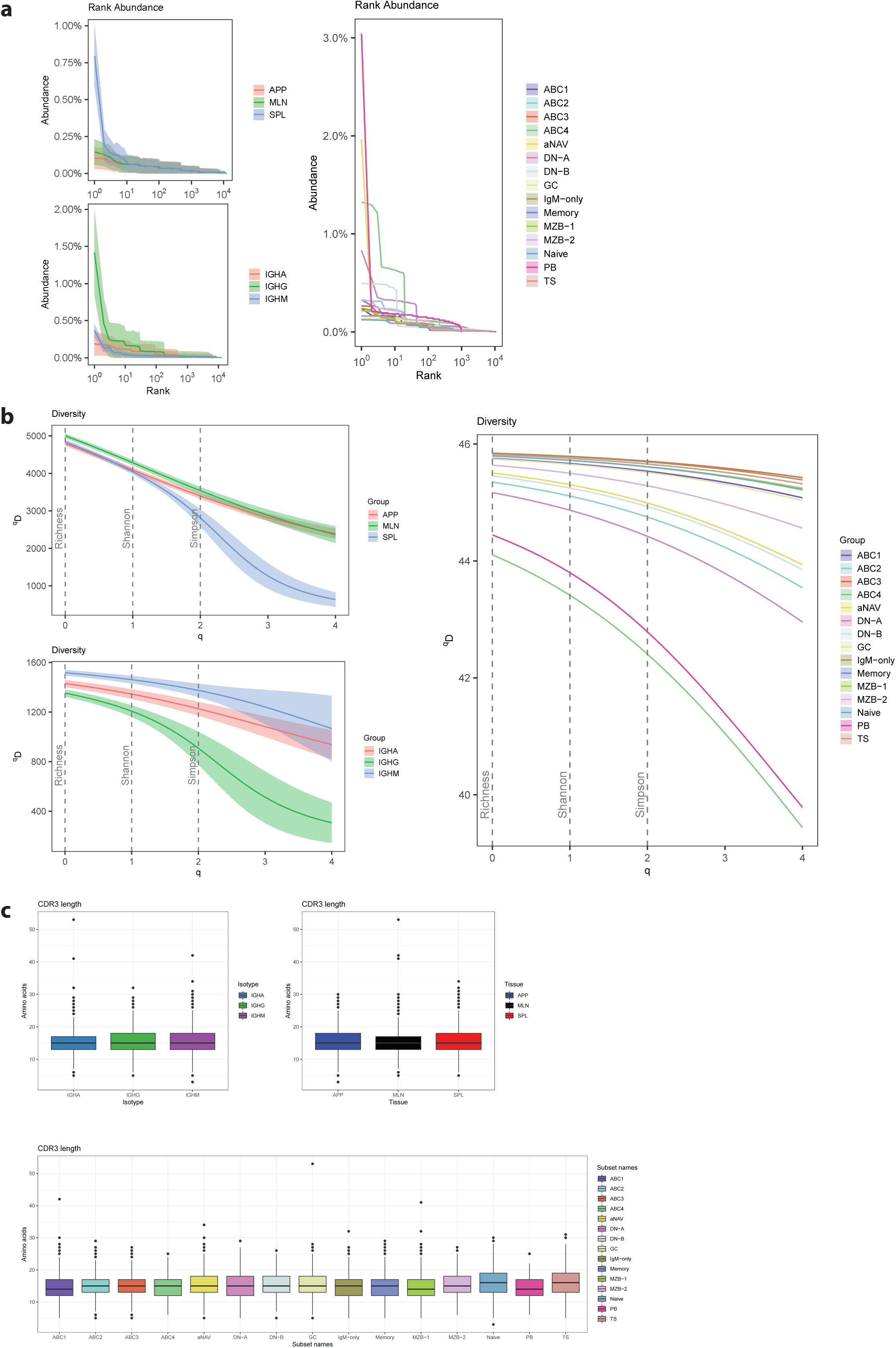
Clonal abundance, diversity, and CDR3 length analysed by tissue, isotype and B cell subset. **(A).** Clone abundance (the size of each clone as a percent of the repertoire) is plotted against the rank of each clone, where the rank is sorted by size from larger (rank 1, left) to smaller (right). The shaded areas are 95% confidence intervals. For visualisation clarity, confidence intervals are not shown for the subsets (no significance). **(B).** Diversity profile curve that plots Hill diversity scores (^q^D) on the y-axis as a function of diversity orders (q). Different values of q represent different measures of diversity: q = 0 (Species Richness), q = 1 (Shannon Entropy), q = 2 (inverse Simpson Index). The shaded areas represent the 95% confidence interval. For visualisation clarity, confidence intervals are not shown for the subsets (no significance). **(C).** Distribution of the CDR3 amino acid length.

**Fig. S6:**
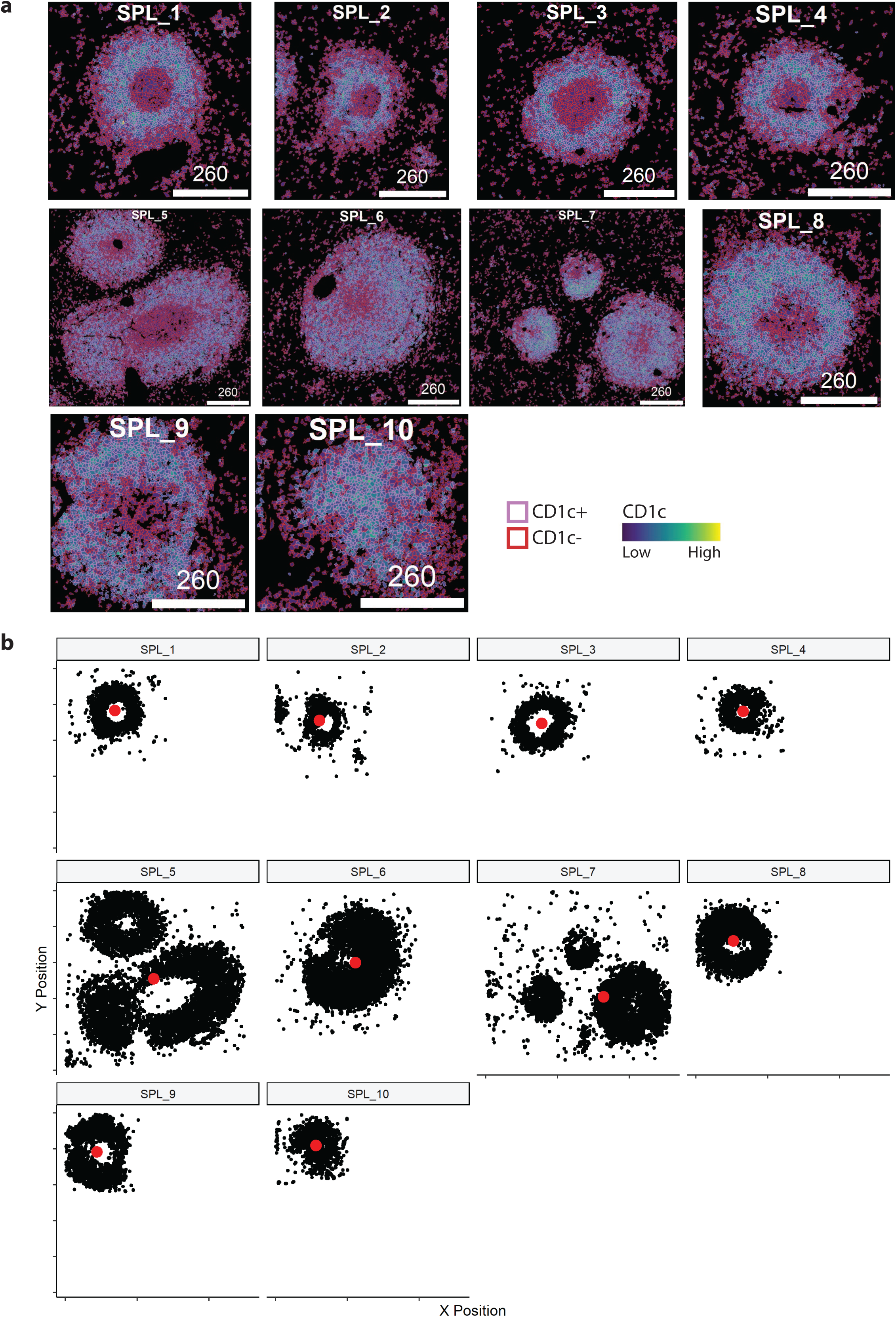
Masking of CD1c^+^ B cells and centre point calculation for image mass cytometry analysis. **(A).** Visualisation of CD1c^+/-^ cell type (coloured by outline) and CD1c marker expression (coloured by fill) on CD20^+^ segmentation masks. **(B).** Scatterplot indicating the location of every CD1c^+^ B cell (black), and the centre point of these cells (red). SPL_5 and SPL_7 were excluded from downstream analysis.

**Fig. S7:**
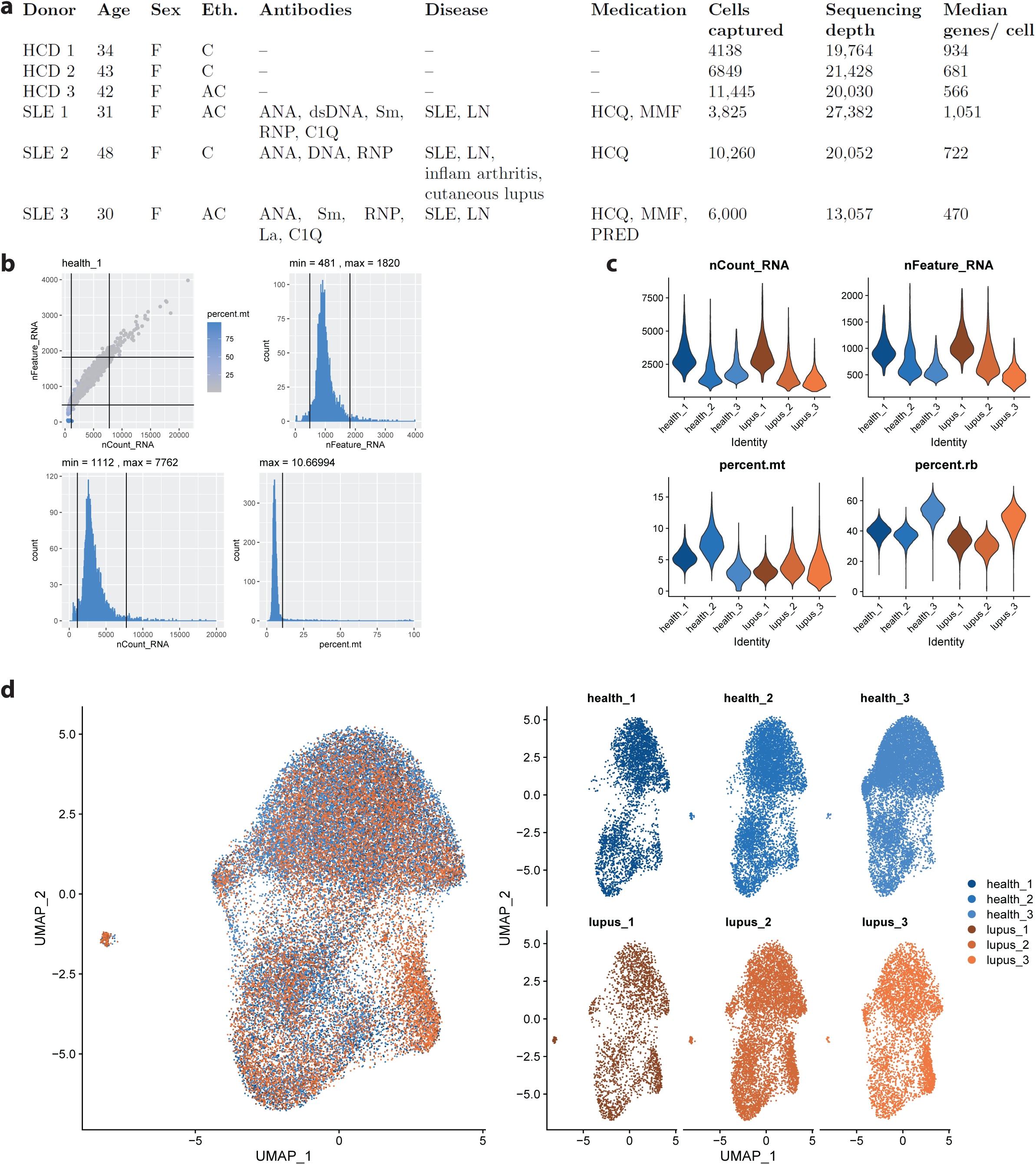
Donor details, 10x genomics workflow, and integration of B cells in blood. **(A).** Demographic details of healthy/SLE patients, cells captured, and sequencing depth. Eth. = Ethnicity, C = Caucasian, AC = African Caribbean, LN = lupus nephritis, MMF = mycophenolate mofetil, PRED = prednisolone, HCQ = hydroxychloroquine. **(B).** Example of quality control processing where thresholds were set automatically based on 3 median absolute deviation. **(C).** Summary of nCount, nFeature, mitochondrial percentage, ribosomal percentage after auto processing. **(D).** UMAP visualisation of B cell composition in blood after integration, coloured by sample.

**Fig. S8:**
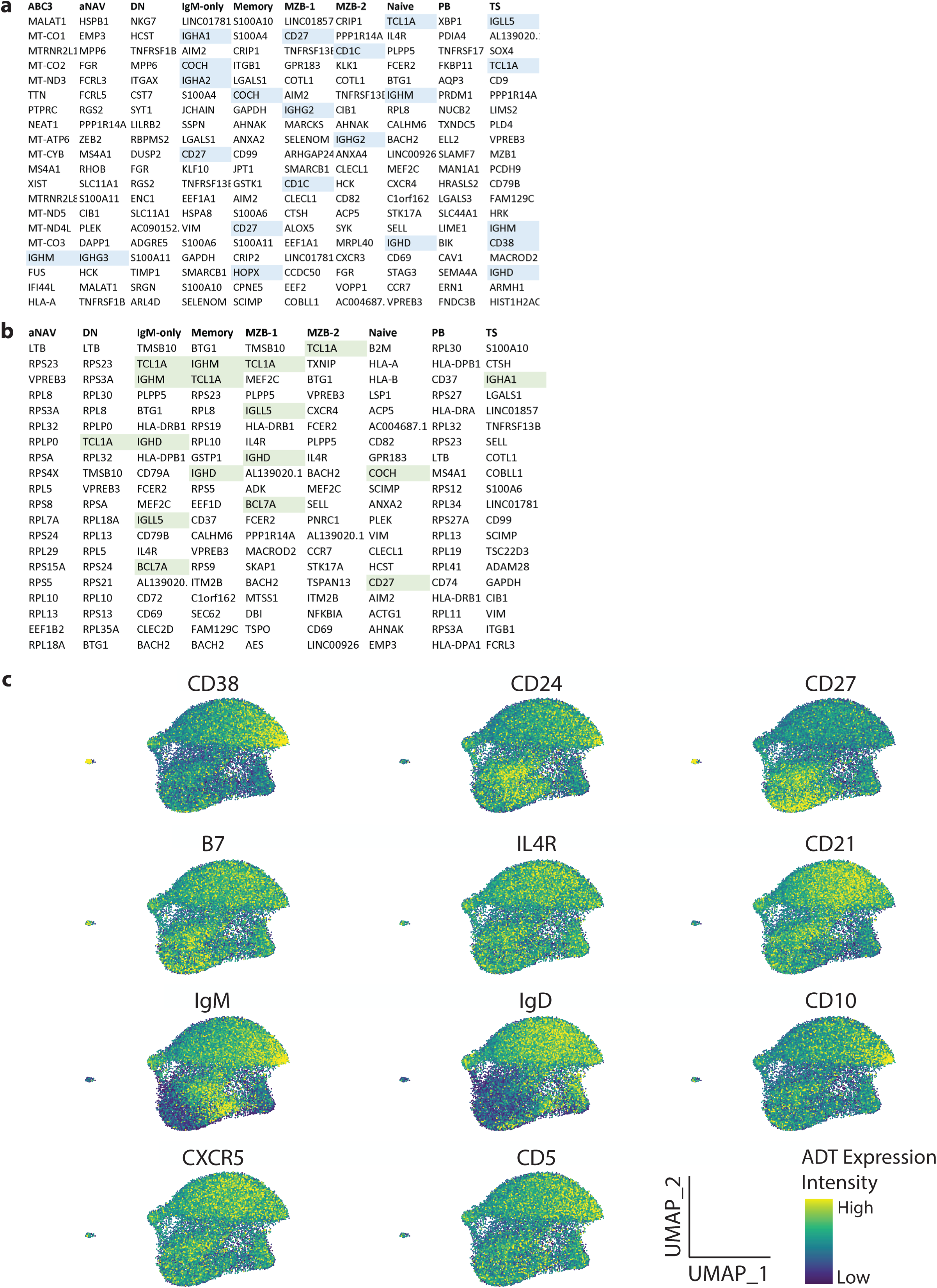
Differentially expressed gene lists for each B cell subtype in blood. **(A).** Top 20 positive differentially expressed genes for each subtype. **(B).** Top 20 negative differentially expressed genes for each subtype. Genes used for annotating cluster are highlighted in blue and green. P values were calculated using Wilcoxon test with Bonferroni correction for multiple comparisons. **(C).** UMAP visualisation of ADT surface protein expression.

**Fig. S9:**
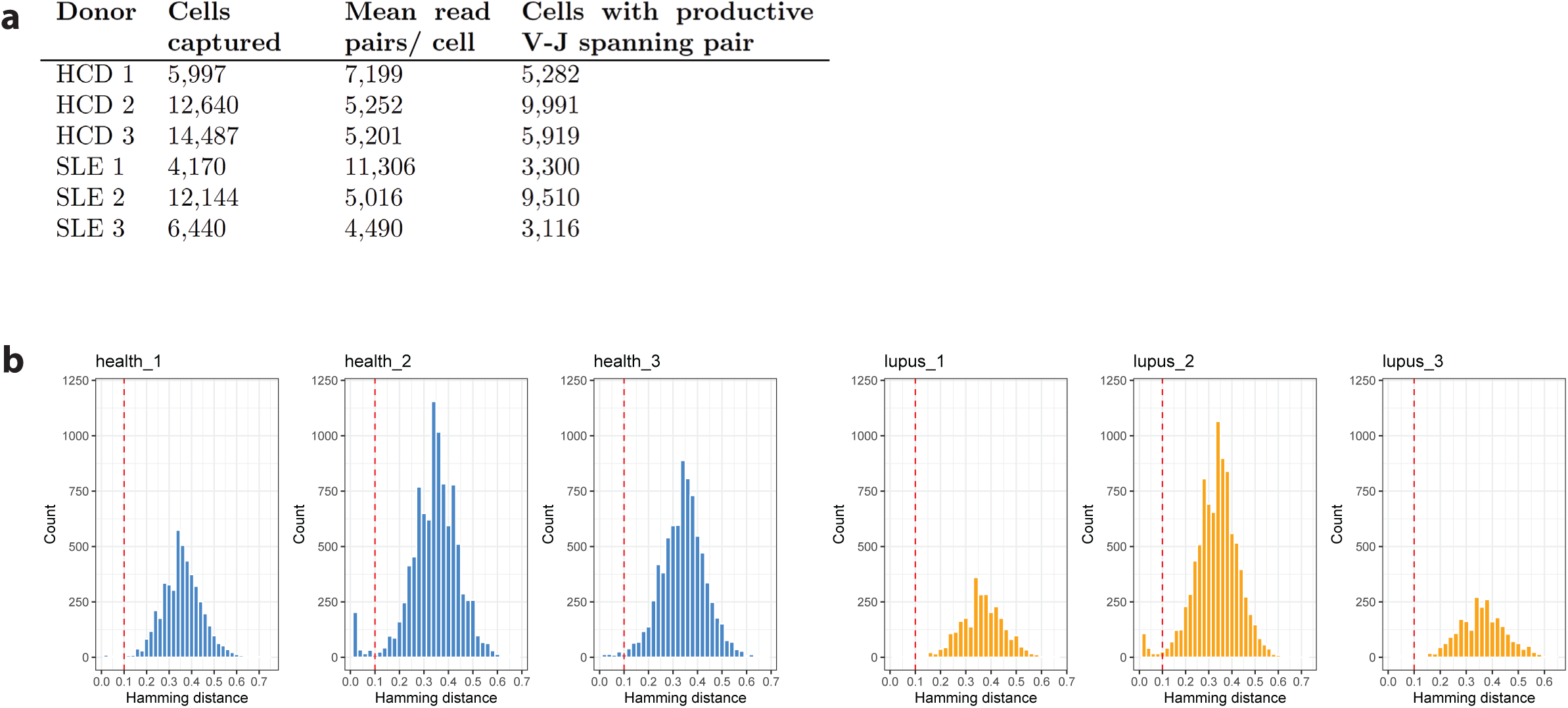
BCR workflow and clonal clustering threshold of blood. **(A).** Cells captured, mean read pairs per cell, number of productive V-J cells for each sample. **(B).** Using the Immcantation pipeline, the distance between sequences and its nearest-neighbour was calculated to determine the threshold for separating clonal groups. The threshold of 0.1 was determined by inspection of the distance-to-nearest plot.

**Supplementary Table 1:** Details of antibodies used for liquid mass cytometry. Metal tags, targets, clone names, working dilutions, and supplier of antibodies used for liquid mass cytometry.

| Metal | Target | Clone | Working Dilution | Supplier |
| --- | --- | --- | --- | --- |
| 158Gb | CD10 | HI10a | 1:200 | Fluidigm |
| 143Nd | CD127 (IL-7R) | A019D5 | 1:200 | Fluidigm |
| 163Dy | CD180 (RP105) | MHR73-11 | 1:200 | Biolegend |
| 163Eu | CD185 (CXCR5) | RF8B2 | 1:200 | Fluidigm |
| 142Nd | CD19 | HIB19 | 1:200 | Fluidigm |
| 159Tb | CD197 (CCR7) | G043H7 | 1:200 | Fluidigm |
| 171Yb | CD20 | 2H7 | 1:200 | Fluidigm |
| 166Er | CD24 | ML5 | 1:100 | Fluidigm |
| 169Tm | CD25 | 2A3 | 1:100 | Fluidigm |
| 164Dy | CD267 (TACI) | 1A1 | 1:200 | Biolegend |
| 155Gd | CD268 (BAFF-R) | 11C1 | 1:200 | Fluidigm |
| 173Yb | CD269 (BCMA) | 19F2 | 1:100 | Biolegend |
| 167Er | CD27 | O323 | 1:200 | Fluidigm |
| 174Yb | CD279 (PD-1) | EH12.2H7 | 1:200 | Fluidigm |
| 154Sm | CD3 | UCHT1 | 1:200 | Fluidigm |
| 170Er | CD307d (FcRL4) | 413D12 | 1:200 | Biolegend |
| 165Ho | CD307e (FcRL5) | 413D12 | 1:200 | Biolegend |
| 144Nd | CD38 | HIT2 | 1:200 | Fluidigm |
| 176Yb | CD4 | RPA-T4 | 1:200 | Fluidigm |
| 162Dy | CD40 | 5C3 | 1:200 | Biolegend |
| 89Y | CD45 | HI30 | 1:200 | Fluidigm |
| 145Nd | CD45RB | MEM-55 | 1:100 | Fluidigm |
| 168Er | CD8 | SK1 | 1:200 | Fluidigm |
| 161Dy | CD80 (B7-1) | 2D10.4 | 1:200 | Fluidigm |
| 148Nd | IgA | Polyclonal | 1:200 | Fluidigm |
| 146Nd | IgD | IA6-2 | 1:200 | Fluidigm |
| 172Yb | IgM | MHM-88 | 1:100 | Fluidigm |

**Supplementary Table 2:** Details of antibodies used for imaging mass cytometry. Metal tags, targets, clone names, working dilutions, and supplier of antibodies used for imaging mass cytometry.

| **Metal** | **Target** | **Clone** | **Working Dilution** | **Supplier** |
| --- | --- | --- | --- | --- |
| 153Eu | DDX21 | EPR14495 | 1:400 | Abcam* |
| 154Sm | IgD | EPR6416 | 1:300 | Abcam* |
| 159Tb | CD68 | KP1 | 1:400 | Fluidigm |
| 161Dy | CD20 | H1 | 1:250 | Fluidigm |
| 164 | CD1c | OTI2F4 | 1:200 | Abcam* |
| 167Er | GzB | EPR20129-217 | 1:300 | Fluidigm |
| 168Er | Ki67 | B56 | 1:400 | Fluidigm |
| 170Er | CD3 | Polyclonal, C-Terminal | 1:800 | Fluidigm |
| 171Yb | CD27 | EPR8569 | 1:300 | Fluidigm |
| 175Lu | CD25 | EPR6452 | 1:50 | Fluidigm |

\* conjugated in-house

